# Nonstructural protein 1 of SARS-CoV-2 is a potent pathogenicity factor redirecting host protein synthesis machinery toward viral RNA

**DOI:** 10.1101/2020.08.09.243451

**Authors:** Shuai Yuan, Lei Peng, Jonathan J. Park, Yingxia Hu, Swapnil C. Devarkar, Matthew B. Dong, Shenping Wu, Sidi Chen, Ivan Lomakin, Yong Xiong

**Author notes:** These authors contributed equally (Shuai Yuan, Lei Peng). Correspondence (S.C.), (I.L.), (Y.X.).

## Abstract

The COVID-19 pandemic affects millions of people worldwide with a rising death toll. The causative agent, severe acute respiratory syndrome coronavirus 2 (SARS-CoV-2), uses its nonstructural protein 1 (Nsp1) to redirect host translation machinery to the viral RNA by binding to the ribosome and suppressing cellular, but not viral, protein synthesis through yet unknown mechanisms. We show here that among all viral proteins, Nsp1 has the largest impact on host viability in the cells of human lung origin. Differential expression analysis of mRNA-seq data revealed that Nsp1 broadly alters the transcriptome in human cells. The changes include repression of major gene clusters in ribosomal RNA processing, translation, mitochondria function, cell cycle and antigen presentation; and induction of factors in transcriptional regulation. We further gained a mechanistic understanding of the Nsp1 function by determining the cryo-EM structure of the Nsp1-40S ribosomal subunit complex, which shows that Nsp1 inhibits translation by plugging the mRNA entry channel of the 40S. We also determined the cryo-EM structure of the 48S preinitiation complex (PIC) formed by Nsp1, 40S, and the cricket paralysis virus (CrPV) internal ribosome entry site (IRES) RNA, which shows that this 48S PIC is nonfunctional due to the incorrect position of the 3’ region of the mRNA. Results presented here elucidate the mechanism of host translation inhibition by SARS-CoV-2, provide insight into viral protein synthesis, and furnish a comprehensive understanding of the impacts from one of the most potent pathogenicity factors of SARS-CoV-2.

**Highlights:** ORF screen identified Nsp1 as a major cellular pathogenicity factor of SARS-CoV-2

Nsp1 broadly alters the gene expression programs in human cells

Nsp1 inhibits translation by blocking mRNA entry channel

Nsp1 prevents physiological conformation of the 48S PIC

## Introduction

SARS-CoV-2, which causes the worldwide COVID-19 pandemic affecting millions of people, belongs to the β-coronaviruses (Coronaviridae Study Group of the International Committee on Taxonomy of, 2020). The virus contains a positive-sense and single-stranded RNA that is composed of 5’-UTR, two large overlapping open reading frames (ORF1a and ORF1b), structural and accessory protein genes, and 3’-poly-adenylated tail (Lim et al., 2016). Upon entering the host cells, ORF1a and ORF1b are translated and proteolytically processed by virus-encoded proteinases to produce functional nonstructural proteins (Nsps) that play important roles in the viral infection and RNA genome replication (Masters, 2006). Nsp1 is the first viral gene encoded by ORF1a (Figure 1A) and is among the first proteins to be expressed after infection (Ziebuhr, 2005). It was shown that human SARS-CoV and group 2 bat coronavirus Nsp1 plays a key role in suppressing the host gene expression (Kamitani et al., 2006; Narayanan et al., 2008; Tohya et al., 2009). SARS-CoV Nsp1 has been shown to inhibit host gene expression using a two-pronged strategy. Nsp1 targets the 40S ribosomal subunit to stall the translation in multiple steps during initiation of translation and also induces an endonucleolytic cleavage of host RNA to accelerate degradation (Kamitani et al., 2009; Lokugamage et al., 2012). Nsp1 therefore has profound inhibitory effects on the host protein production, including suppressing the innate immune system to facilitate the viral replication (Narayanan et al., 2008) and potentially long-term cell viability consequences. Intriguingly, viral mRNA overcomes this inhibition by a yet unknown mechanism, likely mediated by the conserved 5’ UTR region of viral mRNA (Huang et al., 2011; Tanaka et al., 2012). Taken together, Nsp1 acts as an important factor in viral lifecycle and immune evasion, and may be an important virulence factor causing the myriad of long-term illnesses of COVID-19 patients. It has been proposed as a target for live attenuated vaccine development (Wathelet et al., 2007; Zust et al., 2007).

**Figure 1.**
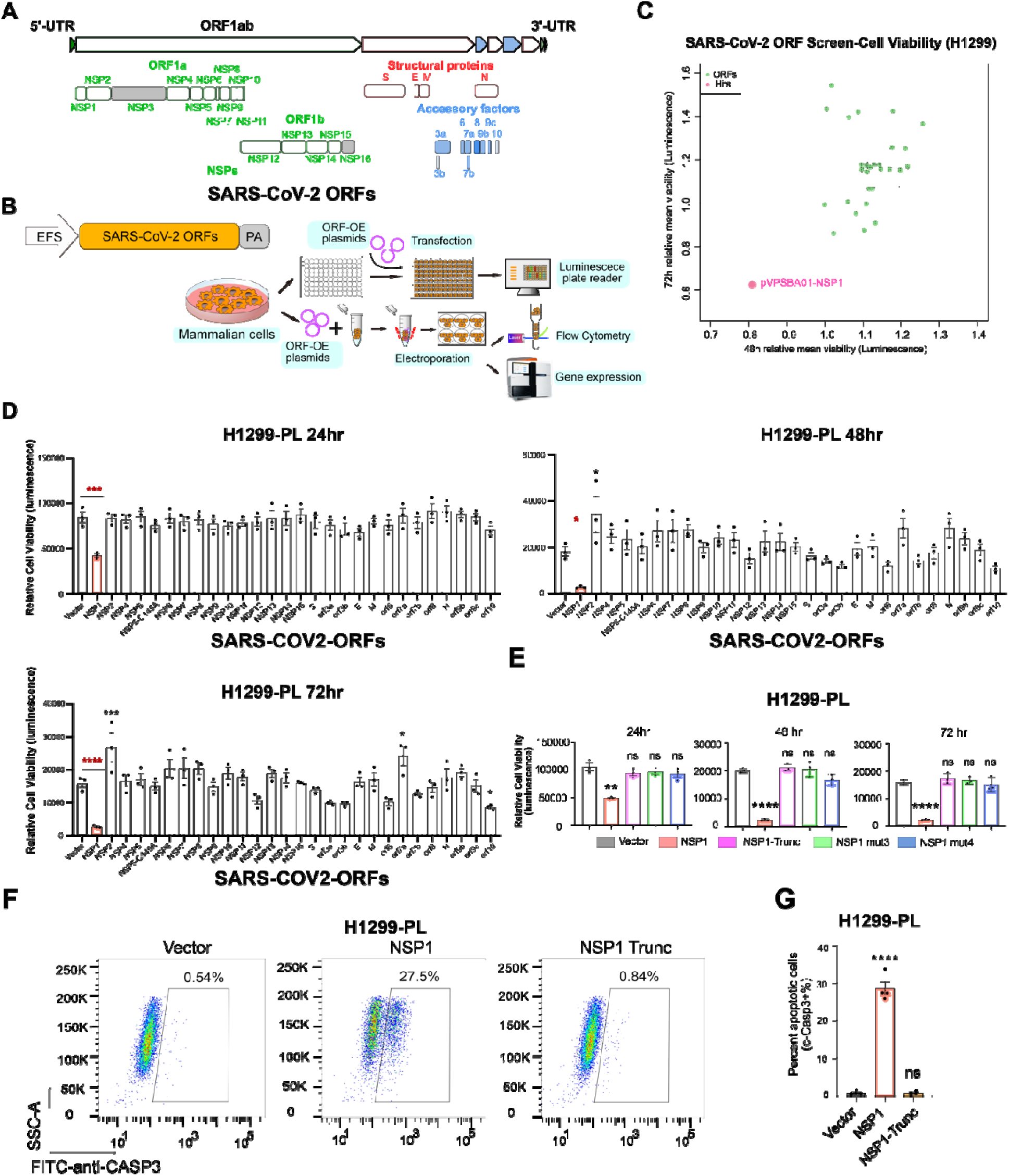
SARS-CoV-2 ORF mini-screen identified Nsp1 as a key viral protein with host cell viability effect. **(A)** Schematics of viral protein coding frames along SARS-CoV-2 genome. Colored ORFs indicate the ones used in this study, while two ORFs in grey are not (Nsp3 and Nsp16). **(B)** Schematics of molecular and cellular experiments of viral proteins. **(C)** Scatter plot of SARS-CoV-2 ORF mini-screen for host viability effect in H1299 cells, at 48 and 72 hours post ORF introduction. Each dot represents the mean normalized relative viability of host cells transfected with a viral protein encoding ORF. Dash line error bars indicate standard deviations. (n = 3 replicates). Pink color indicates hits with p < 0.05 (one-way ANOVA, with multiple group comparison). **(D)** Bar plot of firefly luciferase reporter measurement of viability effects of SARS-CoV-2 ORFs in H1299-PL cells, at 24, 48 and 72 hours post ORF introduction (n= 3 replicates). **(E)** Bar plot of firefly luciferase reporter measurement of viability effects of Nsp1 and three Nsp1 mutants (truncation, mut3: R124S/K125E and mut4: N128S/K129E) in H1299-PL cells, at 24, 48 and 72 hours post ORF introduction (left, middle and right panels, respectively) (n = 3 replicates). **(F)** Flow cytometry plots of apoptosis analysis of Nsp1 and loss-of-function truncation mutant in H1299-PL cells, at 48 hours post ORF introduction. Percentage of apoptotic cells was gated as cleaved Caspase 3 positive cells. **(G)** Quantification of flow-based apoptosis analysis of Nsp1 and loss-of-function truncation mutant in H1299-PL cells, at 48 hours post ORF introduction. For all bar plots in this figure: Bar height represents mean value and error bars indicate standard error of the mean (sem). (n = 3 replicates for each group). Statistical significance was accessed by ordinary one-way ANOVA, with multiple group comparisons where each group was compared to empty vector control, with p-values subjected to multiple-testing correction by FDR method. (ns, not significant; * p < 0.05; ** p < 0.01; *** p < 0.001; **** p < 0.0001). See also Figure S1.

It is common for RNA viruses to target the initiation step of the host protein translation system to allow expression of the viral proteins (Jan et al., 2016). Most cellular mRNAs have a 5’ 7-methylguanosine (m7G) cap structure, which is essential for mRNA recruitment to the 43S preinitiation complex (PIC) through interaction with the translation initiation factor (eIF) eIF4F. 43S PIC is formed by the 40S ribosomal subunit, the ternary complex eIF2-GTP-Met-tRNA^iMet^, and the multi-subunit initiation factor eIF3. Binding of the 43S PIC to the m7G-cap results in the loading of the mRNA in the mRNA-binding channel of the 40S to form the 48S PIC, and scanning of the mRNA from 5’ to 3’ direction under control of eIF1A and eIF1, until the initiation codon AUG is placed in the P site of the 40S. Base pairing of Met-tRNA^iMet^ with AUG results in conformational changes in the 48S PIC for joining the large 60S ribosomal subunit to form the 80S ribosome primed for protein synthesis (Hinnebusch, 2014, 2017b; Hinnebusch et al., 2016). With the exception of the cricket paralysis virus (CrPV), which does not require any host’s eIFs, all other viruses may target different eIFs to redirect the host translational machinery on their own mRNA (Lozano and Martinez-Salas, 2015; Walsh and Mohr, 2011).

We present here data demonstrating that among all viral proteins, Nsp1 causes the most severe viability reduction in the cells of human lung origin. The introduction of Nsp1 in human cells broadly alter the transcriptomes by repressing major gene clusters responsible for protein synthesis, mitochondria function, cell cycle and antigen presentation, while inducing a broad range of factors implicated in transcriptional regulation. We further determined the cryo-EM structures of the Nsp1-40S complex with or without the CrPV IRES RNA, which reveal the mechanism by which Nsp1 inhibits protein synthesis and regulates viral protein production. These results significantly advance our understanding of the Nsp1-induced suppression of host gene expression, the potential mechanisms of SARS-CoV-2 translation initiation, and the broad impact of Nsp1 as a comorbidity-inducing factor.

## Results

### SARS-CoV-2 open reading frame (ORF) screen identifies Nsp1 as a major viral factor that affects cellular viability

A recent study has mapped the interactome of viral protein to host cellular components in human HEK293 cells (Gordon et al., 2020), suggesting that these viral proteins might have diverse ways of interacting or interfering with the fundamental cellular machineries of the host cell. We generated a non-viral over-expression vector (pVPSB) for introduction of viral proteins into mammalian cells and testing their effect on cells (Figure 1B). We first confirmed that the positive control GFP can be introduced into virtually all cells at 100% efficiency, using flow cytometry analysis. We cloned 28 viral proteins (27 of the 29 viral proteins and Nsp5 C145A mutation) as open reading frames (ORFs) into this vector and introduce them into human cells by transfection. We chose to first test H1299, an immortalized cancer cell line of human lung origin. Although H1299 cells are not primary lung epithelial cells, they have been utilized as a cellular model to study SARS-CoV, MERS and SARS-CoV-2 (Hoffmann et al., 2020; Wong et al., 2015).

We introduced all 28 cloned ORFs individually in parallel to conduct a mini-screen of viral proteins’ effect on the viability of H1299 cells (Figures 1B and 1C). We measured cell viability in two time points, 48 and 72 hours (h) post transfection. Unexpectedly, we found Nsp1 as the sole “hit” with significant effect on cell viability at both time points (Figure 1C). To validate the viability observations with increased sensitivity, we generated an H1299 cell line with a constitutive firefly luciferase reporter (H1299-PL), and confirmed that GFP can also be introduced into this cell line at near 100% efficiency (Figures S1A-C). We performed validation experiments, again with all 28 ORFs along with vector control, at 3 different time points (24, 48 and 72h). Across all three time points, Nsp1-transfected H1299 cells have dramatically reduced luciferase signal, an approximation of cell numbers (Figure 1D). We further repeat the same experiments with the Vero E6 cell line, an African monkey (*Cercopithecus Aethiops*) kidney derived cell line, commonly used in SARS-CoV-2 cellular studies (Blanco-Melo et al., 2020; Hoffmann et al., 2020; Kim et al., 2020; Zhou et al., 2020). Consistently, we observed a robust reduction of cellular viability in Vero E6 cells transfected with Nsp1 across all 3 time points (Figure S1D). These data revealed that among all SARS-CoV-2 proteins, Nsp1 has the largest detrimental effect on cell viability in H1299 and Vero E6 cells.

### Nsp1 mutants abolish cellular viability phenotype

To ensure that the observed reduction of cell viability is indeed from expression of functional Nsp1, we tested three different mutants of Nsp1, including a truncation mutation after residues 12 (N terminal mutant) and two double mutations that have been reported to ablate the activity of the highly homologous SARS-CoV Nsp1 (Wathelet et al., 2007). The point mutations include Nsp1 mutant3 that has R124/K125 replaced with S124/E125 (R124S/K125E) and Nsp1 mutant4 that has N128/ K129 replaced with S128/E129 (N128S/K129E). We performed cellular viability assays with wild-type (WT) Nsp1 along with all three of its mutants. In both H1299-PL and Vero E6-PL cells, we again observed that introduction of Nsp1 into cells significantly reduced cell viability along 24, 48, and 72 hours post electroporation (Figures 1E and S1E). Each of the three mutants (truncation, R124S/K125E and N128S/K129E) reverted this phenotype to the vector control level, fully abolishing the cytotoxic effect of Nsp1 (Figures 1E and S1E). These results confirmed that functional Nsp1, but not its loss-of-function mutants, induce reduction of cellular viability when overexpressed in the two mammalian cell lines.

We further tested if Nsp1 expression also leads to cell death. We introduced Nsp1 into H1299 cells, along with controls of empty vector and several other viral proteins (Nsp2, Nsp12, Nsp13, Nsp14, ORF9b, and Spike), and measured cellular apoptosis at 48h post electroporation by flow cytometry analysis of cleaved Caspase 3 staining. We found that introduction of Nsp1, but not other viral proteins, induced apoptosis in H1299 cells (Figure S1F). To ensure the cellular apoptosis effect is indeed from expression of functional Nsp1 protein, we performed the same apoptosis assay with Nsp1 and the three non-functional mutants described above. Consistently, only wild-type (WT) Nsp1 induced apoptosis in H1299-PL cells, whereas the three mutants did not (Figure S1G). Replicates of this cleaved Caspase 3 flow assay with the truncation mutation of Nsp1 confirmed that WT Nsp1, but not the loss-of-function truncation mutant, induced apoptosis in H1299-PL cells (Figures 1F and 1G).

### Transcriptome profiling of Nsp1-overexpressed cells

To unbiasedly investigate the global gene expression changes induced by Nsp1 or its loss-of-function mutant form, we performed transcriptome profiling. We first confirmed that Nsp1 is indeed over-expressed in host cells by qPCR using a custom-designed NSP1-specific probe, at both 24 and 48 hours post electroporation (Figure 2A). We then electroporated in quadruplicates for each of Nsp1, its truncation mutant, or vector control plasmid into H1299-PL cells, and collected samples 24 hours post electroporation for mRNA-seq. We collected 24h instead of 48h or 72h samples in order to capture the earlier effect of Nsp1 on cellular transcriptome. We mapped the mRNA-seq reads to the human transcriptome and quantified the expression levels of annotated human transcripts and genes (Table S3). Principle component analysis showed clear grouping and separation of WT Nsp1, mutant Nsp1, or vector control groups (Figure 2B), confirming the overall quality of the Nsp1 mRNA-seq dataset.

**Figure 2.**
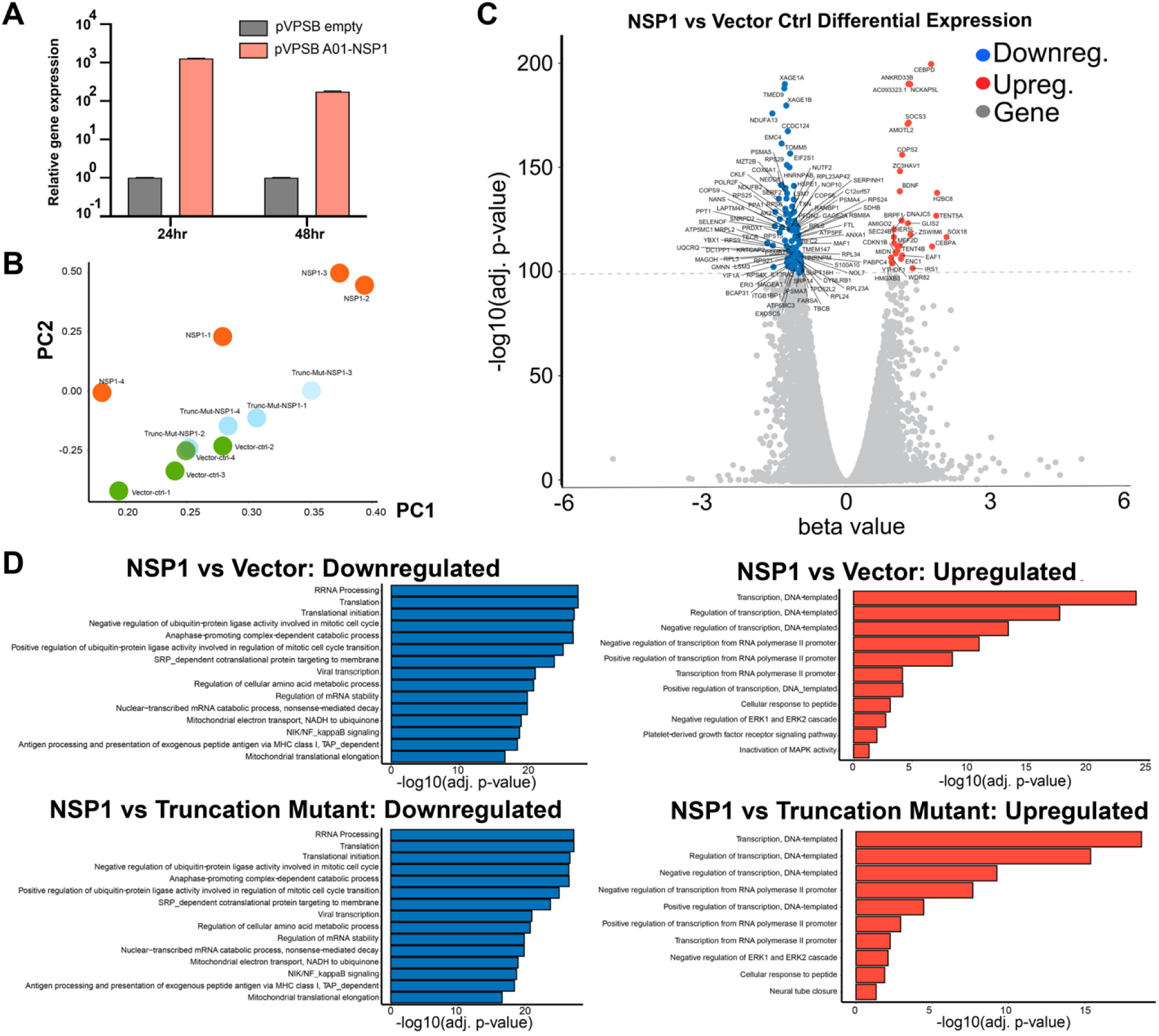
Transcriptome profiling of H1299 cells introduced with NSP1 and NSP1 truncation mutant by RNA-seq. **(A)** Quantitative PCR (qPCR) confirmation of *NSP1* overexpression, at 24 and 48 hours post electroporation. (n = 3 replicates). **(B)** Principle component analysis (PCA) plot of the entire mRNA-seq dataset, showing separation between Nsp1, Vector control and Nsp1 truncation mutant groups, all electroporated into H1299-PL cells and harvested 24 hours post electroporation. RNA samples were collected as quadruplicates (n = 4 each group). **(C)** Volcano plot of differential expression between of Nsp1 vs Vector Control electroporated cells. Top differentially expressed genes (FDR adjusted q value < 1e-100) are shown with gene names. Upregulated genes are shown in orange. Downregulated genes are shown in blue. **(D)** Bar plot of top enriched pathway analysis by DAVID Biological Processes (BP). Nsp1 vs Vector control (top), or Nsp1 vs Nsp1 mutant (top), highly downregulated (left) and upregulated (right) genes are shown (q < 1e-30). See also Figure S2

Differential expression analysis revealed broad and potent gene expression program changes induced by Nsp1 (Figure 2C; Table S3 and S4), with 5,394 genes significantly downregulated and 3,868 genes significantly upregulated (FDR adjusted q value < 0.01). To examine the highly differentially expressed genes, we used a highly stringent criteria (FDR adjusted q value < 1e-30), and identified 1,245 highly significantly downregulated genes (top NSP1 repressed genes) and 464 highly significantly upregulated genes (top Nsp1 induced genes) (Figure 2C; Table S3 and S4). In sharp contrast, Nsp1 truncation mutant and the vector control showed no differential expression in the transcriptome, even when using the least stringent criteria (FDR adjusted q value < 0.05) (Figures S2A-B; Table S3 and S4). These data revealed that Nsp1 alone can cause major alterations broadly in the transcriptome shortly (24h) after its introduction into host cells, consistent with its cell viability phenotype (Figure 1).

### Enriched pathway analysis on differentially expressed gene sets revealed strong signatures of cellular transcriptome alterations by Nsp1

We globally examined the highly differentially expressed genes as a result of Nsp1 expression. To understand what these genes represent as a group, we performed DAVID clustering and biological processes (BP) analysis on the 1,245 top Nsp1-repressed genes and the 464 top Nsp1-induced genes, respectively (Figure 2D; Table S4). Enriched pathways in the top Nsp1-repressed genes showed that the most significant gene ontology groups include functional annotation clusters of ribosomal proteins and translation related processes, such as terms of ribonucleoprotein (RNP) (Hypergeometric test, FDR-adjusted q = 6.30e-57), ribosomal RNA processing (q = 2.03e-28), and translation (q = 3.93e-28). Highly enriched Nsp1-repressed genes also include the clusters of mitochondria function and metabolism (most terms with q < 1e-15) and cell cycle and cell division (most terms with q < 1e-10), consistent with the reduced cell viability phenotype. Other intriguing enriched Nsp1-repressed pathways include ubiquitin/proteasome pathways and antigen-presentation activities, as well as mRNA processing. We further performed gene set enrichment analysis (GSEA) that takes into consideration both gene set and ranks of enrichment, and the results largely validated the DAVID findings, with highly similar strongly enriched pathways (Figures 3A and S2C). Analysis of highly differentially expressed genes between Nsp1 vs. Nsp1 mutant showed results virtually identical to those of Nsp1 vs. vector (Figures S2A-B, Table S4).

**Figure 3.**
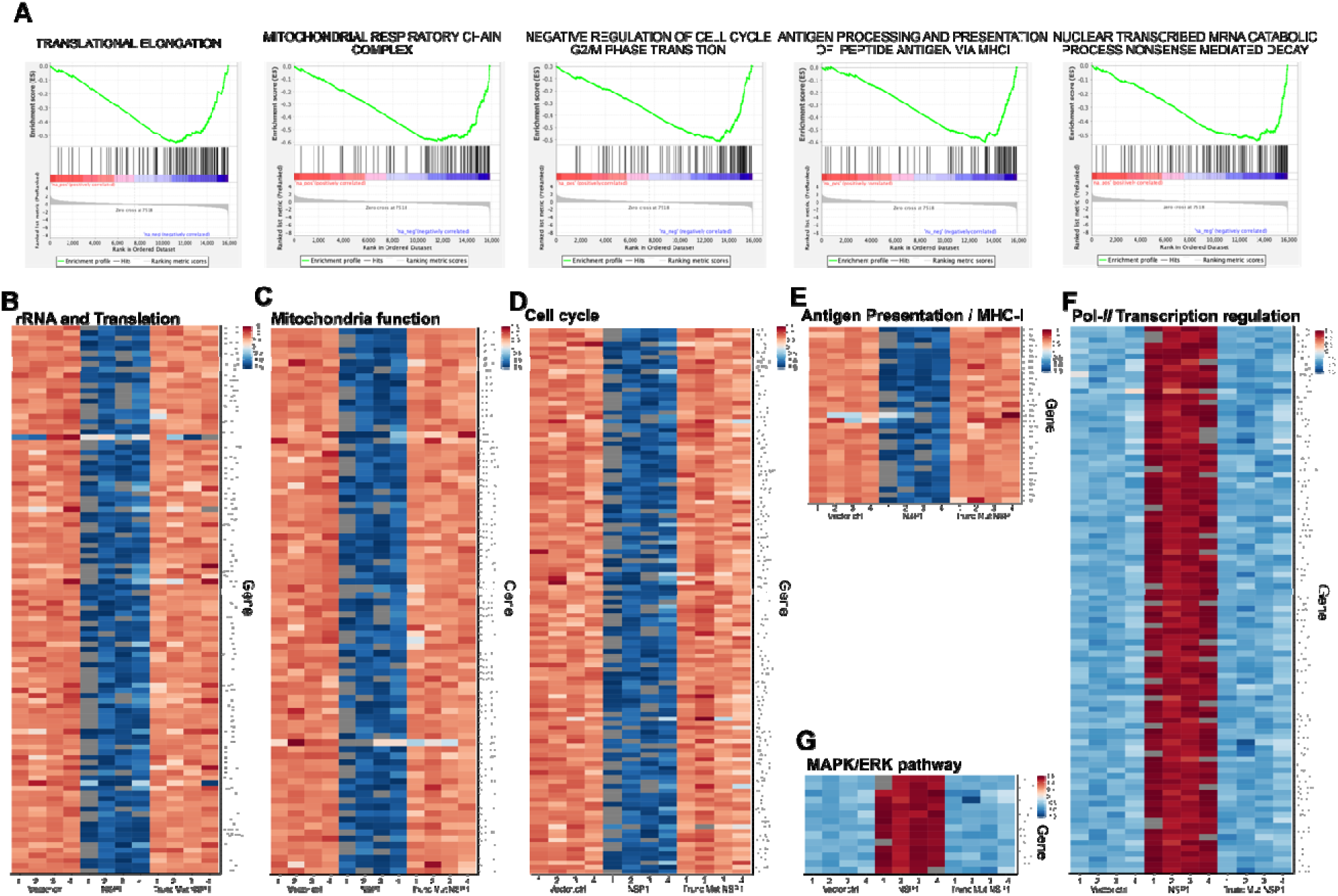
Highly differentially expressed genes between Nsp1, Vector control and Nsp1 mutant group in the context of top major enriched pathways. **(A)** Gene set enrichment plots of representative enriched pathways by GSEA. **(B-E)** Heatmap of Nsp1 highly repressed genes (q < 1e-30) in rRNA processing and translation (**B**), mitochondria function (**C**), cell cycle (**D**), MHC-I antigen presentation processes (**E**). **(F-G)** Heatmap of Nsp1 highly induced genes (q < 1e-30) in *polII* related transcription regulation processes (**F**) and the MAPK/ERK pathway (**G**). See also Figure S2

We then examined the expression levels of the highly differentially expressed genes in the context of enriched pathways in Nsp1, mutant Nsp1, or vector control plasmid in H1299-PL cells. As shown in the heatmaps (Figure 3B), over 70 genes involved in translation are strongly repressed upon introduction of Nsp1, including the RPS, RPL, MRPS, MRPL family members, along with other translational regulators such as *AKT1*. The repression effect on these genes is completely absent in the Nsp1 mutant group (Figure 3B). The strong repression effect also hit multiple members of the gene families involved in mitochondria function, such as the COX, NUDFA, NUDFB and NUDFS families (Figure 3C). Consistent with the cellular phenotypes, Nsp1 also repressed a large number of mitotic cell cycle genes, including members in the CDK, CDC and CCNB families, components of the centrosome, the anaphase promoting complex and various kinases (Figure 3D). While part of the signal may be driven by ribosomal and/or proteosomal genes, multiple genes involved in the mRNA processing and/or nonsense-mediated decay nevertheless are significantly repressed by Nsp1(Figures S2D-E). Interestingly, DAVID BP enrichment analysis of Nsp1-repressed genes also scored the antigen presentation pathway, mostly proteasome components along with several MHC-I component members (Figure 3E). Concordantly, Nsp1-repressed genes are also enriched in the ubiquitination and proteasome degradation pathways (Figure S2F).

On the other hand, genes highly induced by Nsp1 hit a broad range of factors implicated in transcriptional regulation, such as unfolded protein response regulators (*ATF4, XBP1*), FOX family transcription factors (TFs) (*FOXK2, FOXE1, FOXO1, FOXO3*), Zinc finger protein genes (*ZFN217*, *ZFN567*), KLF family members (*KLF2, KLF10*), SOX family members (*SOX2, SOX4*), Homeobox genes (*HOXD9, HOXC8, HOXD13*), GATA TFs (*GATAD2B, GATA6*), dead-box protein genes (*DDX5, DHX36*), cell fate regulators (*RUNX2, CREBRF, LIF, JUNB, ELK1, JAG1, SMAD7, BCL3, EOMES*); along with certain epigenetic regulators of gene expression such as the SWI/SNF family members *ARID1A, ARID1B, ARID3B*, and *ARID5B* (Figure 3F). Interestingly, highly upregulated genes are also slightly enriched in the MAPK/ERK pathway, where Nsp1 expression induces multiple DUSP family members (Figure 3G). The upregulated genes also include several KLF family members related to the process of cellular response to peptide (Figure S2G). Again, the induction effect on these genes is completely abolished in the Nsp1 mutant group (Figures 3F and 3G). These data together showed that Nsp1 expression broadly and significantly altered multiple gene expression programs in the host H1299-PL cells.

### Cryo-EM structure reveals Nsp1 is poised to block host mRNA translation

To elucidate the mechanism of translation inhibition by Nsp1, we determined the cryo-EM structure of rabbit 40S ribosomal subunit complex with Nsp1 at 2.7 Å resolution (Table 1, Figures S3 and S4). The quality of the cryo-EM map allowed us to unambiguously identify Nsp1 that binds to the head and the body domains of the 40S around the entry to the mRNA channel (Figure 4). The density observed in the mRNA entry channel enabled us to build an atomic model for the C-terminal domain of Nsp1 (C-Nsp1, amino acids (aa) 145-180) (Figure 4A). C-Nsp1 comprises two α-helices (α1, aa 154-160; α2, aa 166-179) and two short loops (aa 145-153 and 161-165), which blocks the mRNA entry channel (Figure 4B). Besides the α-helices in the mRNA channel, extra globular density between the ribosomal protein uS3 and rRNA helix h16 is observed at a lower contour level, whose dimensions roughly matched the N-terminal domain of Nsp1 (aa: 13-127, N-Nsp1, PDB:2HSX) (Almeida et al., 2007) (Figure 4C). However, N-Nsp1 does not appear to be stably bound to the 40S and the low local resolution of the cryo-EM map in this region did not allow for an atomic model for the N-Nsp1.

**Figure 4.**
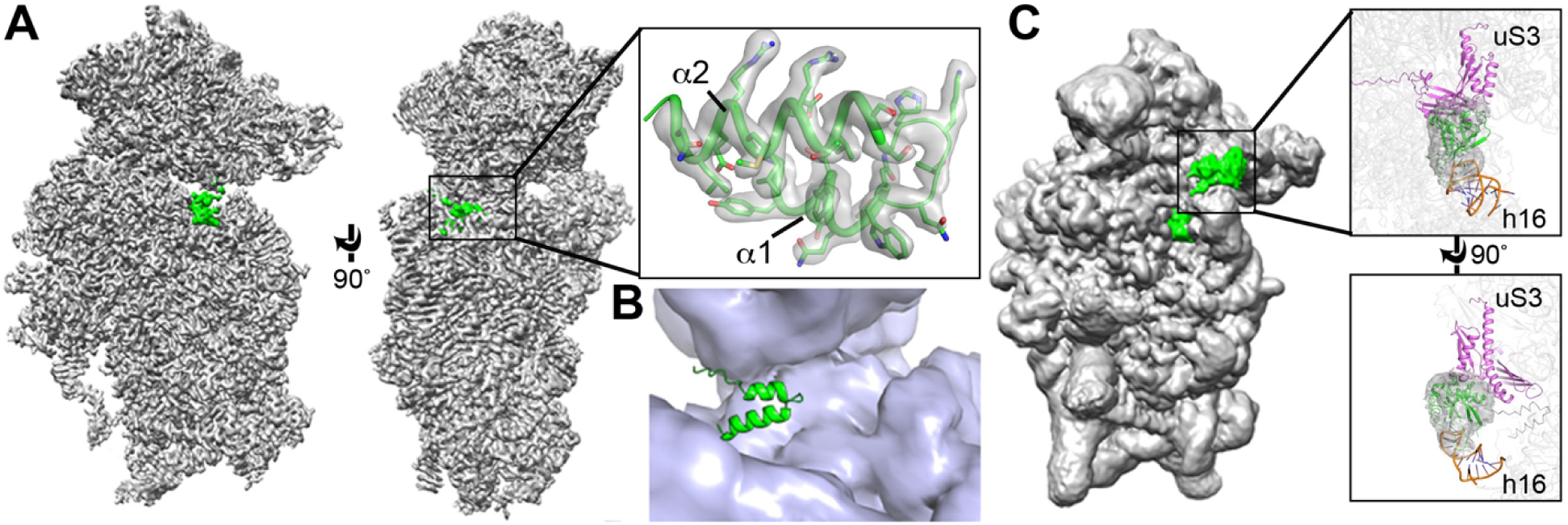
cryo-EM structure of the Nsp1-40S ribosome complex. **(A)** Overall density of the Nsp1-40S ribosome complex with Nsp1 (green) and 40S. ribosome (gray). Inset shows C-Nsp1 with corresponding density with clear sidechain features. C-Nsp1 α-helices (α1, aa 154-160; α2, aa 166-179) are labeled. **(B)** Cross section of the C-Nsp1 (green) within the mRNA entry channel. 40S. ribosome is shown in surface and C-Nsp1 is displayed in cartoon. **(C)** Overall density of Nsp1-40S ribosome complex at a lower contour level. Insets. shows the extra globular density with SARS-CoV Nsp1 N-terminal domain (PDB:2HSX, green) fitted. Ribosomal protein uS3 (magenta) and rRNA h16 (orange) are shown in cartoon. See also Figures S3 and S4.

**Table 1.**
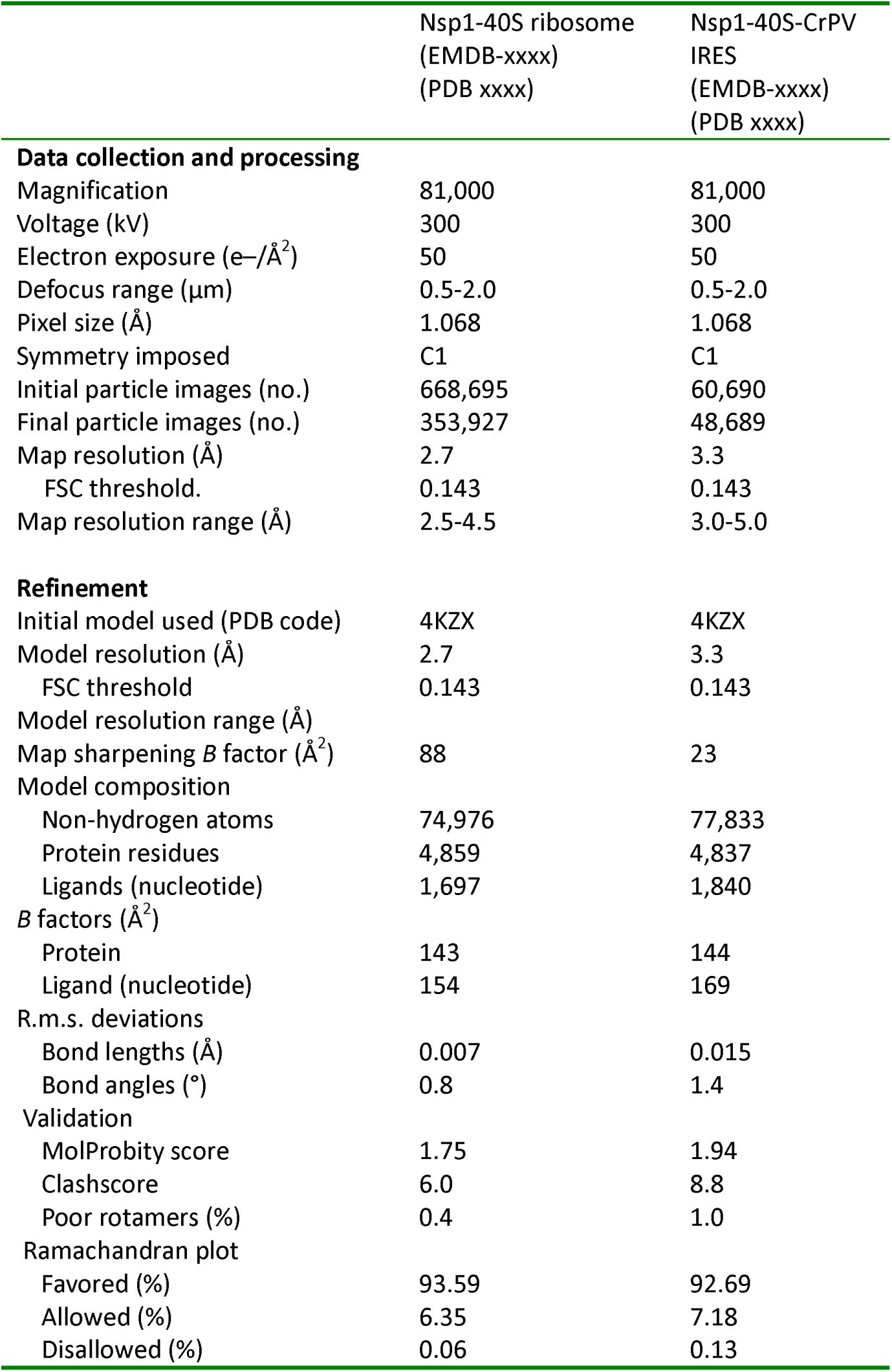
Cryo-EM data collection, refinement and validation statistics

|  | Nsp1-40S ribosome<br>(EMDB-xxxx)<br>(PDB xxxx) | Nsp1-40S-CrPV<br>IRES<br>(EMDB-xxxx)<br>(PDB xxxx) |
| --- | --- | --- |
| **Data collection and processing** |  |  |
| Magnification | 81,000 | 81,000 |
| Voltage (kV) | 300 | 300 |
| Electron exposure (e-/Å<sup>2</sup>) | 50 | 50 |
| Defocus range (μm) | 0.5-2.0 | 0.5-2.0 |
| Pixel size (Å) | 1.068 | 1.068 |
| Symmetry imposed | C1 | C1 |
| Initial particle images (no.) | 668,695 | 60,690 |
| Final particle images (no.) | 353,927 | 48,689 |
| Map resolution (Å) | 2.7 | 3.3 |
| FSC threshold. | 0.143 | 0.143 |
| Map resolution range (Å) | 2.5-4.5 | 3.0-5.0 |
| **Refinement** |  |  |
| Initial model used (PDB code) | 4KZX | 4KZX |
| Model resolution (Å) | 2.7 | 3.3 |
| FSC threshold | 0.143 | 0.143 |
| Model resolution range (Å) |  |  |
| Map sharpening B factor (Å<sup>2</sup>) | 88 | 23 |
| Model composition |  |  |
| Non-hydrogen atoms | 74,976 | 77,833 |
| Protein residues | 4,859 | 4,837 |
| Ligands (nucleotide) | 1,697 | 1,840 |
| B factors (Å<sup>2</sup>) |  |  |
| Protein | 143 | 144 |
| Ligand (nucleotide) | 154 | 169 |
| R.m.s. deviations |  |  |
| Bond lengths (Å) | 0.007 | 0.015 |
| Bond angles (°) | 0.8 | 1.4 |
| Validation |  |  |
| MolProbity score | 1.75 | 1.94 |
| Clashscore | 6.0 | 8.8 |
| Poor rotamers (%) | 0.4 | 1.0 |
| Ramachandran plot |  |  |
| Favored (%) | 93.59 | 92.69 |
| Allowed (%) | 6.35 | 7.18 |
| Disallowed (%) | 0.06 | 0.13 |

C-Nsp1 bridges the head and body domains of the 40S ribosomal subunit through extensive electrostatic and hydrophobic interactions with the ribosomal proteins uS3 of the head, uS5 and eS30 and helix h18 of the 18S rRNA in the body (Figure 5A). The buried surface area of interaction between C-Nsp1 and the 40S ribosomal subunit is ∼1,420 Å^2^. The negatively charged residues D152, E155 and E159 of C-Nsp1 interact with the positively charged residues R117, R116, R143 and K148 of uS3, respectively (Figure 5B). In addition, the positively charged surface of C-Nsp1 binds to the negatively charged rRNA backbone of h18 (Figure 5C). K164 of Nsp1 inserts into the negatively charged pocket formed by the backbone of G625 and U630 of the rRNA h18. H165 of Nsp1 stacks with the base of U607 of h18, and R171 and R175 of C-Nsp1 interact with the negatively charged patch formed by G601, A604, G606 and U607 of h18 (Figure 5C). Besides electrostatic contacts, a large hydrophobic patch of C-Nsp1, which is formed by F157, W161, L173 and L177, interacts with a complimentary hydrophobic patch on uS5 formed by V106, I109, P111, T122, F124, V147 and I151 (Figure 5D). Intriguingly, K164 and H165 of Nsp1, which have been shown to play an important role in host translation inhibition, are conserved only in the betacoronaviruses (beta-CoVs) (Figure 5E). In addition, the other Nsp1 residues interacting with the h18 of rRNA are also conserved only among the beta-CoVs (Figure 5E). This sequence conservation indicates that the hydrophobic interactions between C-Nsp1 and uS5 are likely universal in both alpha-and beta-CoVs, while the electrostatic interactions between C-Nsp1 and the h18 of the 18S rRNA are conserved only in the beta-CoVs.

**Figure 5.**
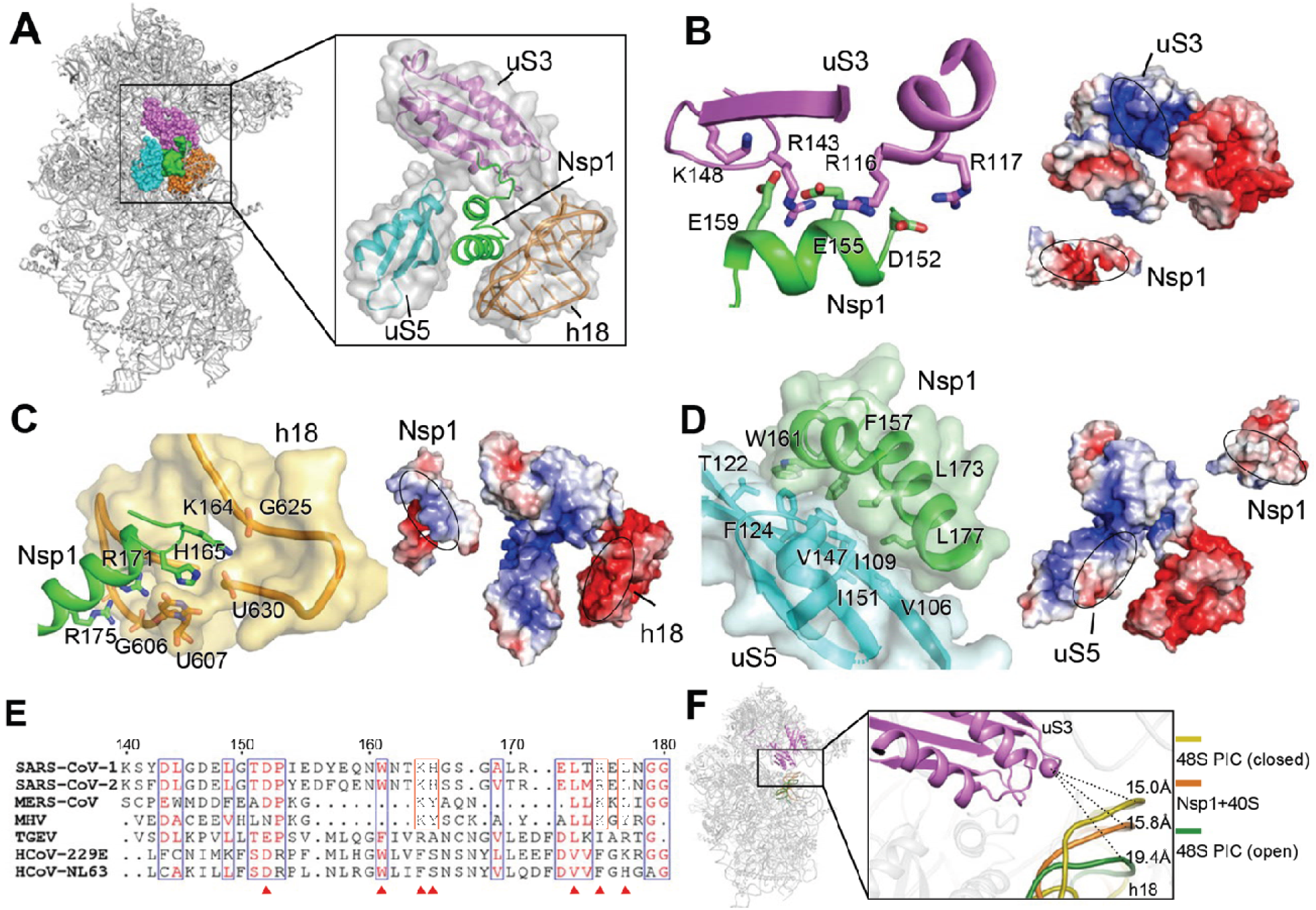
Structural basis of C-Nsp1 and 40S ribosome interaction. **(A)** Overall structure of the C-Nsp1-40S ribosome complex, with C-Nsp1 (green. surface) and the surrounding protein uS3 (magenta sphere representation), uS5 (cyan) and rRNA h18 (orange) highlighted. The inset shows zoomed-in view of C-Nsp1 in cartoon, with the surrounding 40S components in cartoon and surface to illustrate the mRNA entry channel. **(B-D)** Molecular interactions between C-Nsp1 and 40S ribosome components, including uS3 **(B)**, h18 **(C)**, uS5 **(D)**. Left panels: Proteins and rRNA are in the same color as in **(A)** and shown in cartoon, with binding pocket and hydrophobic interface depicted in surface. The interacting residues are shown in sticks. Right panels: The complementary electrostatic surfaces at the interfaces (marked with ovals), colored by electrostatic potential (blue, positively charged; red, negatively charged). **(E)** Alignment of the last 40 residues at Nsp1 C-terminus from beta-CoVs (SARS-CoV-1, SARS-CoV-2, MERS-CoV and MHV) and alpha-CoVs (TGEV, HCoV-229E and HCoV-NL63) coronaviruses. Residues conserved in both alpha- and beta-CoVs are boxed in blue. Residues only conserved in beta-CoVs coronaviruses are with orange boxes. Conserved residues that mediate the interaction with the 40S are marked with red triangles. **(F)** The conformation of the 40S ribosome in the Nsp1-40S complex is similar to the. close form in the 48S PIC. Q179 of uS3 (magenta cartoon) is displayed as a sphere. h18 is in cartoon and colored dark yellow (48S closed conformation), orange (Nsp1-40S ribosome complex) and dark green (48S open conformation), with distances to Q179 indicated by the dashes.

The extensive interactions result in C-Nsp1 plugging the mRNA entry channel, which prevents the loading and accommodation of the mRNA (Figure 4B), providing a structural basis for the inhibition of host protein synthesis by Nsp1 of SARS-CoV-2 and SARS-CoV reported previously (Kamitani et al., 2009; Kamitani et al., 2006). Because Nsp1 molecules of both viruses share 84% amino acid sequence identity, they likely act by the same mechanism (Figures 5A and 5E). It was shown that K164 and H165 of SARS-CoV Nsp1 KH motif are essential for the suppression of host protein synthesis (Kamitani et al., 2009). In our structure the motif provides critical interactions with helix h18, anchoring Nsp1 to the 18S rRNA (Figure 5C). These interactions constitute ∼15% of the overall C-Nsp1-40S ribosome interacting surface, which explains the detrimental effect of K164A and H165A mutations on inhibition of host protein synthesis.

### Nsp1 locks the 40S in a conformation incompatible with mRNA loading and disrupts initiation factor binding

The ribosomal protein uS3 is conserved in all kingdoms. Together with h16, h18 and h34 of 18S rRNA it constitutes the mRNA-binding channel and the mRNA entry site (Graifer et al., 2014; Hinnebusch, 2017a). It has been shown that uS3 interacts with the mRNA and regulates scanning-independent translation on a specific set of mRNAs (Haimov et al., 2017; Sharifulin et al., 2015). Interestingly, conserved residues R116 and R117 of uS3, which are crucial for stabilizing mRNA in the entry channel and maintaining 48S PIC in the closed conformation, are interacting with D152, E155 of Nsp1 in our structure (Dong et al., 2017; Hinnebusch, 2017a) (Figure 5B). Moreover, the conformation of the 40S ribosomal subunit in Nsp1-40S complex is similar to that of ‘closed state’ of 48S PIC with initiator tRNA locked in the P site and the latch closed (Lomakin and Steitz, 2013), which is incapable of mRNA loading. The distance between G610 (h18) and GLN179 (CA, uS3) is shortened from 19.4 Å in the ‘open state’ 48S PIC (PDB:3JAQ) to 15.8 Å in Nsp1-40S ribosomal complex, which is similar to the distance of 15.0 Å in the closed state 48S PIC (PDB:4KZZ) (Figure 5F). This shows that Nsp1 not only plugs the mRNA entry channel, but also keeps the 40S subunit in a conformation that is incompatible with mRNA loading.

The known structure of the N-terminal domain of SARS-CoV (N-Nsp1) (Almeida et al., 2007) (PDB ID: 2HSX) can be docked into the extra globular density between uS3 and rRNA helix h16 in the cryo-EM map (Figure 6A). This potential interaction between N-Nsp1 and uS3 covers most of the uS3 surface on the solvent side, including the GEKG loop of uS3 (aa: 60-63) that corresponds to the consensus GXXG loop conserved in the KH domains of various RNA-binding proteins (Babaylova et al., 2019b; Graifer et al., 2014). Mutation of the GEKG loop to alanines does not abrogate the ability of the 40S to bind mRNA and form 48S preinitiation complex (PIC). Instead, it results in the formation of aberrant 48S PIC that cannot join the 60S ribosomal subunit and assemble the 80S initiation complex (Graifer et al., 2014). Peculiarly, binding of SARS-CoV Nsp1 to the ribosome led to the same effect (Kamitani et al., 2009). We hypothesize that Nsp1 may prevent the formation of physiological conformation of the 48S PIC induced by uS3 interaction with translation initiation factors, such as the j subunit (eIF3j) of the multi-subunit initiation factor eIF3 (Babaylova et al., 2019b; Cate, 2017; Sharifulin et al., 2016). The eIF3 complex plays a central role in the formation of the translation initiation complex (Cate, 2017; Hinnebusch, 2014). eIF3j alone binds to the 40S ribosomal subunit and stabilizes the interaction with eIF3 complex (Fraser et al., 2004b; Sokabe and Fraser, 2014). The binding site of eIF3j to 40S subunit is not precisely determined. Cryo-EM and biochemical studies mapped it onto the mRNA binding channel of the 40S, extending from the decoding center toward the mRNA entry region, including the GEKG loop of uS3 (Aylett et al., 2015; Fraser et al., 2007; Hershey, 2015) (Figure 6B).

**Figure 6.**
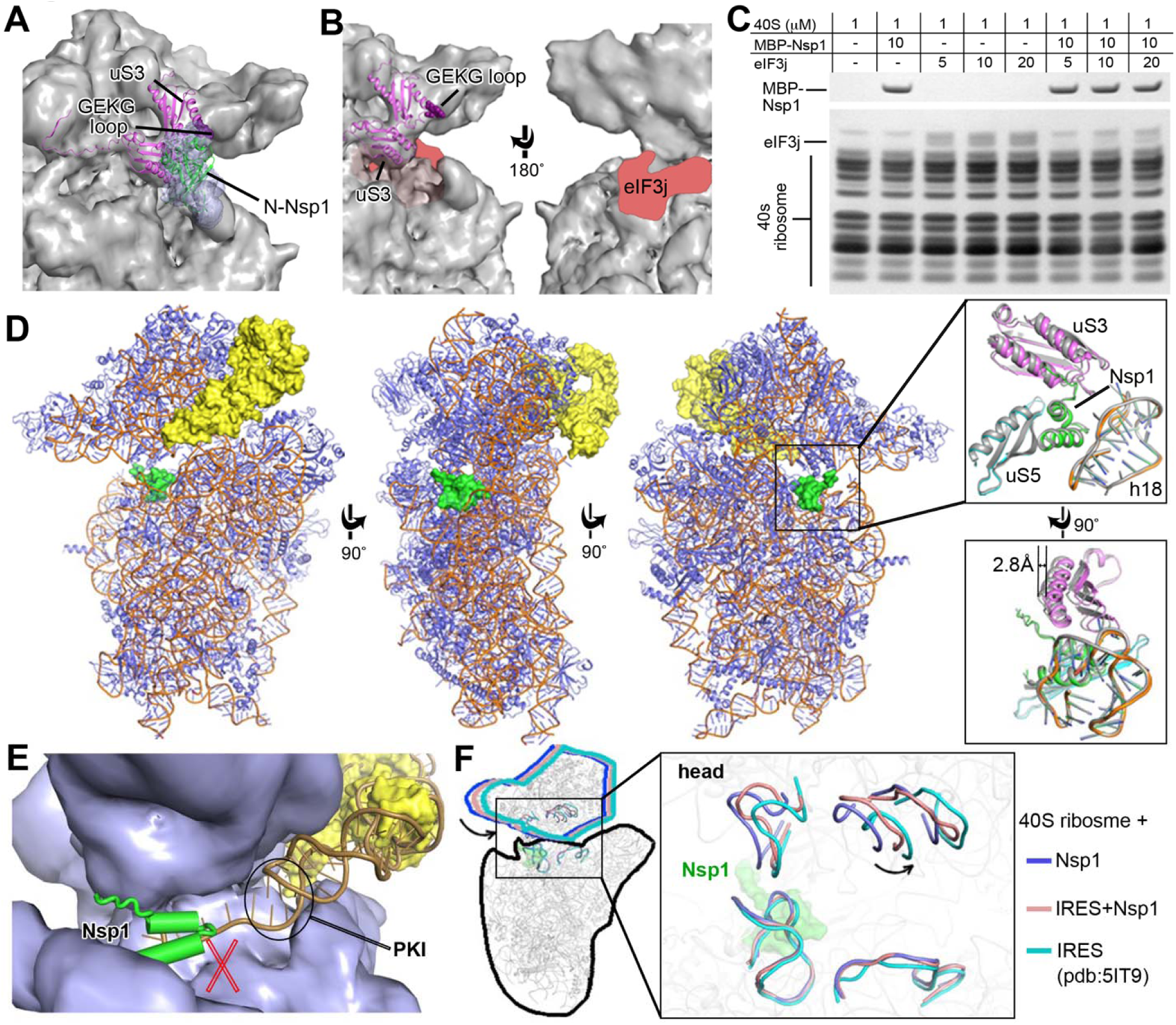
Nsp1 disrupts initiation factor binding and prevents physiological conformation of the 48S PIC. **(A)** The N-terminal domain of Nsp1 covers uS3 surface on the solvent side. The. cryo-EM density in this region is shown in blue surface with SARS-CoV Nsp1 N-terminal domain (PDB:2HSX) fitted. uS3 (magenta) is depicted cartoon. The GEKG loop (dark purple) is shown in sphere representation. **(B)** Potential binding region of eIF3j. The putative location of eIF3j is marked in red. **(C)** SDS-PAGE analysis of Nsp1 and eIF3j competition at different concentration ratios (indicated in the top table). **(D)** Overall structure of the Nsp1-40S-CrPV IRES complex. Nsp1 (green) and IRES. (yellow) are presented in surface. The ribosome proteins (slate) and rRNA (orange) are shown in cartoon. The right insets display the conformation change in the Nsp1-binding region (cartoon representation) with or without the IRES. **(E)** The previously reported model of CrPV IRES (PDB: 5IT9, orange cartoon) fitted. to 40S ribosome in the present of Nsp1 (green cartoon). 40S ribosome (slate) and the currently observed IRES (yellow) are presented in surface. **(F)** C-Nsp1 restricts the 40S ribosome head rotation. Superposition of the Nsp1-40S, Nsp1-40S-CrPV IRES and IRES-40S (PDB:5IT9) complexes is shown is cartoon. Zoomed view displays the head rotations represented by selected rRNA regions. C-Nsp1 (green) is displayed in surface. See also Figures S5, S6 and S7.

We tested if Nsp1 can compete with eIF3j for the binding to the 40S ribosomal subunit. The result showed that Nsp1 indeed significantly reduces the binding between eIF3j and the 40S (Figure 6C). The binding competition of eIF3j and Nsp1 to the 40S was tested at different concentrations. There is little eIF3j binding to the 40S when the concentration of eIF3j is equal or lower than that of Nsp1, and residual eIF3j binding was observed only when its concentration is higher than that of Nsp1 (Figures 6C and S5). By contrast, the binding of Nsp1 to the 40S is not affected even when eIF3j is in excess. These results indicate that Nsp1 disrupts the binding of eIF3j to the 40S, potentially by shielding the access to uS3 and the mRNA binding channel and/or by making the conformation of the 40S unfavorable for eIF3j interaction.

### Nsp1 prevents physiological conformation of the 48S PIC

It was shown previously that binding of SARS-CoV Nsp1 to the 40S ribosomal subunit does not inhibit 48S PIC formation, but it suppresses 60S subunit joining (Kamitani et al., 2009). To understand the effect of Nsp1 of SARS-CoV-2 on 48S PIC, we determined a 3.3 Å resolution cryo-EM structure of Nsp1 bound to the 48S PIC assembled with the cricket paralysis virus (CrPV) internal ribosome entry site (IRES) (Figures 6D, S6, and S7). CrPV IRES has become an important model for studies of the eukaryotic ribosome during initiation, as it is able to directly recruit and assemble with 40S or 80S ribosome without requiring any eIFs (Martinez-Salas et al., 2018). It was shown that SARS-CoV Nsp1 inhibits translation of the CrPV IRES RNA (Kamitani et al., 2009). We first examined whether Nsp1 affects binding of the IRES RNA to the 40S ribosomal subunit. The result shows that Nsp1 and CrPV IRES can bind 40S ribosomal subunit simultaneously (Figure S6). Consistently, both C-Nsp1 and the CrPV IRES can be seen in the cryo-EM map (Figure 6D), where the Nsp1 C-terminal domain is inserted in the RNA entry channel in the same way as in the Nsp1-40S complex without the IRES RNA (Figures 4A and 4B). The local environment of C-Nsp1 in the ribosome RNA entry channel with or without the IRES RNA is quite similar. No conformational changes were observed for C-Nsp1, protein uS5 and rRNA h18, however, the head of the 40S subunit is moved by about 2.8 Å (Figure 6D) (discussed more below).

We fitted the high resolution structure of the CrPV IRES from the yeast 40S-CrPV IRES complex(Murray et al., 2016) (PDB: 5IT9) into our cryo-EM map. Importantly, the pseudoknot I (PKI) domain of the CrPV IRES, which is a structural mimic of the canonical tRNA-mRNA interaction, is not seen in the cryo-EM map, suggesting that it is dislodged from the 40S in the presence of Nsp1 (Figure 6E). Consistently, there would be a clash between Nsp1 C-terminal domain and the 3’ region of the IRES RNA in the previously observed conformation bound to the 40S (Murray et al., 2016) (Figure 6E). The conformation of the 40S head in the Nsp1-40S-CrPV IRES complex is different from that in the Nsp1-40S complex (Figure 6F). The head in the Nsp1-40S-CrPV IRES complex is in somewhat intermediate conformation compared to the Nsp1-40S and the 40S-CrPV IRES complexes (Figure 6F). This suggests that the Nsp1-40S interactions resist the conformational changes induced by the IRES for translation initiation. Conformational changes of the head domain of the 40S subunit play important role in the mRNA loading and recruitment of the 60S subunit to form the 80S ribosome. Nsp1 limits the rotation of the head, which may have profound consequences interfering with the joining of the 60S subunit and the formation of the 80S initiation complex.

## Discussion

SARS-CoV-2 infection causes a series of damages to the human body, often leading to long-term illnesses (Grasselli et al., 2020). However, the cellular phenotypes and the relative contributions of individual viral proteins are not clearly understood. While viral infection is a complex process involving multiple components, certain viral proteins are often in high abundance in cells during active viral replication (Astuti and Ysrafil, 2020; Yoshimoto, 2020). Therefore, understanding the effects of each individual viral protein on the cells provides important insights on the cellular impacts of viral infection. Using a reductionist approach, we tested the gross cellular effect of expressing all the SARS-CoV-2 proteins individually. Among all 27 viral proteins, Nsp1 showed the strongest deleterious effect on cell viability in H1299 cells of human lung epithelial origin. This is in concordance with previous observations from related coronaviruses, such as mouse hepatitis virus (MHV) Nsp1 being a major pathogenicity factor strongly reducing cellular gene expression (Zust et al., 2007), and SARS-CoV Nsp1 inhibiting interferon (IFN)-dependent signaling and having significant effects on cell cycle (Wathelet et al., 2007). A recent study shows that SARS-CoV-2 Nsp1 shuts down mRNA translation in cells and suppresses innate immunity genes such as *IFNb* and *IL-8*, although these experiments were conducted in HEK293T cells of kidney origin, and only a small number of host genes were tested (Thoms et al., 2020). As an unbiased interrogation of global cellular pathways affected by Nsp1, our transcriptome profiling data and gene set enrichment analysis revealed strong signatures of transcriptomic changes in broad ranges of host genes with several major clusters, providing a comprehensive understanding of the impacts of one of the most potent pathogenicity protein factors of SARS-CoV-2 in human cells of lung origin.

Our structure of the SARS-CoV-2 Nsp1 protein bound to the 40S ribosomal subunit establishes a mechanistic basis of the cellular effects of Nsp1, revealing a multifaceted mechanism of inhibition of the host protein synthesis at the initiation stage by the virus. Nsp1 plugs the mRNA channel entry from the position +10 and up, which physically blocks access to the channel by any mRNA (Figure 4B). This is consistent with the result obtained from similar structural studies (Thoms et al., 2020). Moreover, Nsp1 interacts with the ribosomal protein uS3 of the head domain and uS5 of the body domain of the 40S subunit as well as with the helix h18 of the 18S rRNA, which locks the head domain of the 40S subunit in the closed position. This position is characterized by the closed conformation of the “mRNA entry channel latch” that clams around incoming mRNA (Hinnebusch, 2017b; Lomakin and Steitz, 2013; Passmore et al., 2007). The latch is supposed to be closed during the scanning of the mRNA, keeping mRNA locked in the binding cleft and increasing processivity of the scanning, whereas the open conformation of the latch would facilitate the initial attachment of the 43S PIC to the mRNA (Lomakin and Steitz, 2013). Therefore, when Nsp1 keeps the latch closed it makes impossible for the host mRNA to be loaded. In addition, the N-terminal domain of Nsp1 interacts with the KH-domain of uS3, specifically with its GEKG loop crucial for translation initiation (Figures 6A and 6B) (Babaylova et al., 2019a). We showed that Nsp1 competes with eIF3j for the binding to the 40S subunit (Figure 6C). This allows us to propose that Nsp1 weakens the binding of the eIF3 to the 40S subunit by disrupting uS3-eIF3j interaction. Moreover, accessibility to the GEKG loop of uS3 is required for the functional 48S PIC formation (Babaylova et al., 2019a; Fraser et al., 2004a; Graifer et al., 2014; Sokabe and Fraser, 2014).

Our results explain how Nsp1 inhibits protein synthesis; however, how SARS-CoV-2 escapes this inhibition and initiate translation of its own RNA still remains unanswered. The 5’-UTR of SARS-CoV-2 is essential for escaping Nsp1-mediated suppression of translation (Tanaka et al., 2012). Interactions involving the viral 5’ UTR presumably result in the “unplugging” of Nsp1 from the 40S ribosome during the initiation of viral translation. In addition, the weakening of eIF3 binding to the 40S subunit is beneficial for translation initiation of some viruses. The hepatitis C virus (HCV) IRES displaces eIF3 from the interface of the 40S subunit to load its RNA in the mRNA binding channel (Hashem et al., 2013; Niepmann and Gerresheim, 2020). HCV IRES interacts with eIF3a, eIF3c and other core subunits of eIF3 to promote formation of the viral 48S PIC (Cate, 2017). The eIF3d subunit of the eIF3 complex can be cross-linked to the mRNA in the exit channel of the 48S PIC, it has its own cap-binding activity which can replace canonical eIF4E dependent pathway and promote translation of selected cellular mRNAs (Lee et al., 2016; Pisarev et al., 2008; Walker et al., 2020). Interestingly, a recent genome-wide CRISPR screen revealed the eIF3a and eIF3d are essential for SARS-CoV-2 infection (Wei et al., 2020). It is possible that SARS-CoV-2 may use an “IRES-like” mechanism involving eIF3 recruitment by 5’ UTR to overcome Nsp1 inhibition. Binding of 5’ UTR may cause conformational change of the 40S head leading to the latch opening, Nsp1 dissociation, viral RNA loading into mRNA binding channel and formation of the functional 80S initiation complex primed for viral protein synthesis. However, the detailed mechanisms of viral escape of Nsp1 inhibition must await for future experimental studies.

## Acknowledgements

We thank the Yale cryo-EM facilities for assistance with data collection. We thank the Xiong lab and Chen lab members for discussions. We thank Xiaoyun Dai, Lupeng Ye and several others for sharing various plasmids and reagents. We thank the Yale Center for Genome Analysis and Center for Research Computing for providing High-throughput sequencing and computing assistance and resources.

## Author contributions

S.Y., L.P., S.C., I.L. and Y.X. initiated the project and designed the experiments. S.Y. I.L, and Y.H. produced proteins and 40S ribosomal subunit. S.Y. and S.D. performed binding assays. S.Y. prepared the cryo-EM samples. Y.H. and S.W. carried out cryo-EM data collection. S.Y. and Y.X. did cryo-EM data processing. S.Y., I.L., S.D. and Y.X. analyzed cryo-EM structure. L.P. and M.B.D. performed cellar assays. L.P. and J.J.P. performed and processed mRNA-seq. S.Y., L.P., S.C., I.L. and Y.X. prepared the manuscript. S.C., I.L. and Y.X. jointly supervised the work.

## Supplemental Figures

**Figure S1.**
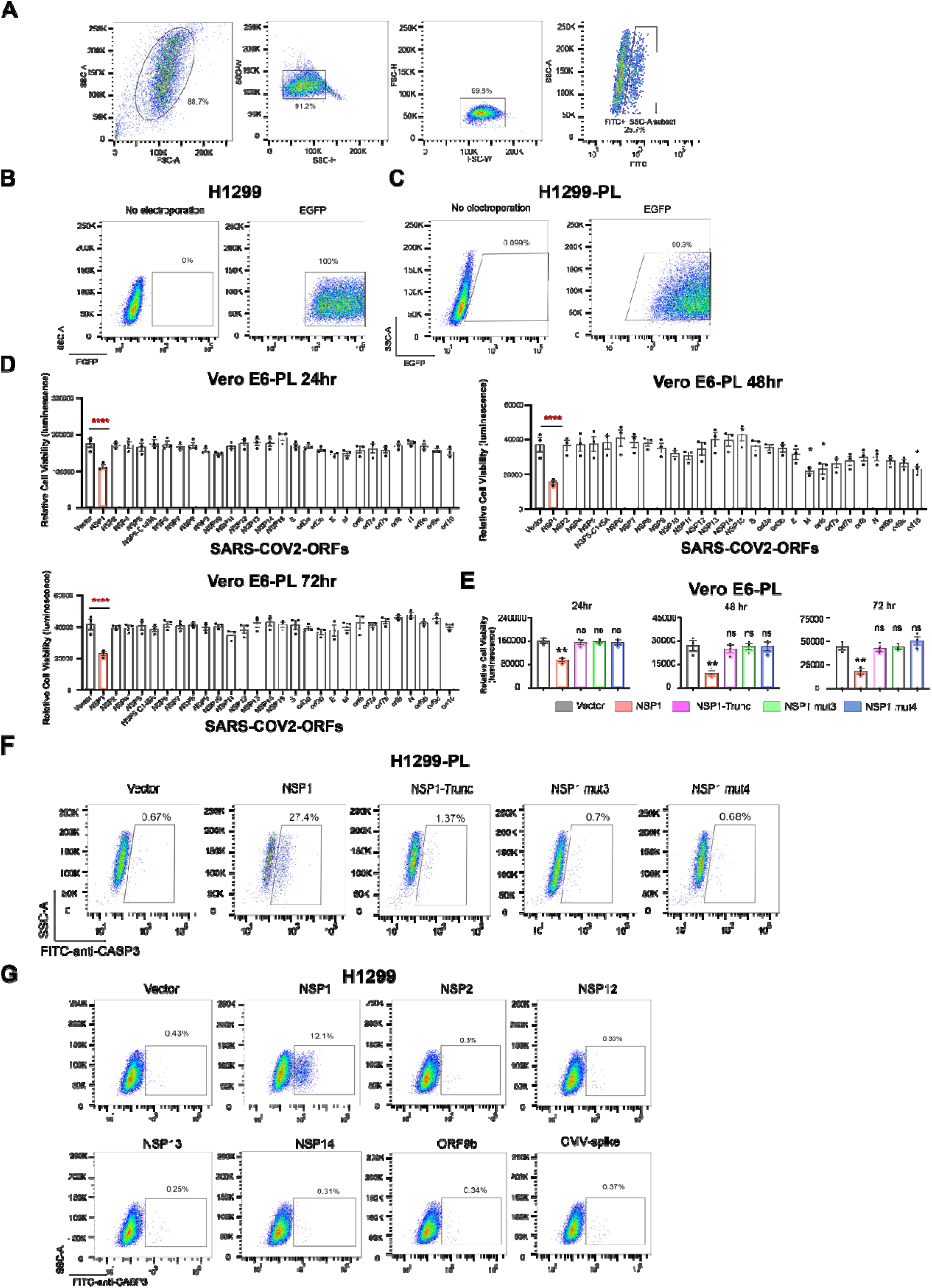
Flow cytometry analysis of cellular effects of SARS-CoV-2 ORFs, Nsp1, and Nsp1 mutants, Related to Figure 1. **(A)** Diagram of example flow gating. **(B)** Flow cytometry plots of GFP expression in H1299 cells, at 48 hours post ORF introduction. **(C)** Flow cytometry plots of GFP expression in H1299-PL cells, at 48 hours post ORF introduction. **(D)** Bar plot of firefly luciferase reporter measurement of viability effects of SARS-CoV-2 ORFs in Vero E6-PL cells, at 24, 48 and 72 hours post ORF introduction (n = 3 replicates). **(E)** Bar plot of firefly luciferase reporter measurement of viability effects of Nsp1 and three Nsp1 mutants (truncation, mut3: R124S/K125E and mut4: N128S/K129E) in Vero E6-PL cells, at 24, 48 and 72 hours post ORF introduction (left, middle and right panels, respectively) (n = 3 replicates). **(F)** Flow cytometry plots of apoptosis analysis of Nsp1 and three Nsp1 mutants (truncation, mut3: R124S/K125E and mut4: N128S/K129E) in H1299-PL cells, at 48 hours post ORF introduction. Percentage of apoptotic cells was gated as cleaved Caspase 3 positive cells. **(G)** Flow cytometry plots of apoptosis analysis of several SARS-CoV-2 ORFs (Nsp1, Nsp2, Nsp12, Nsp13, Nsp14, Orf9b and Spike), at 48 hours post ORF introduction, in H1299 cells. Percentage of apoptotic cells was gated as cleaved Caspase 3 positive cells. For all bar plots in this figure: Bar height represents mean value and error bars indicate standard error of the mean (sem). (n = 3 replicates for each group). Statistical significance was accessed by ordinary one-way ANOVA, with multiple group comparisons where each group was compared to empty vector control, with p-values subjected to multiple-testing correction by FDR method. (ns, not significant; * p < 0.05; ** p < 0.01; *** p < 0.001; **** p < 0.0001).

**Figure S2.**
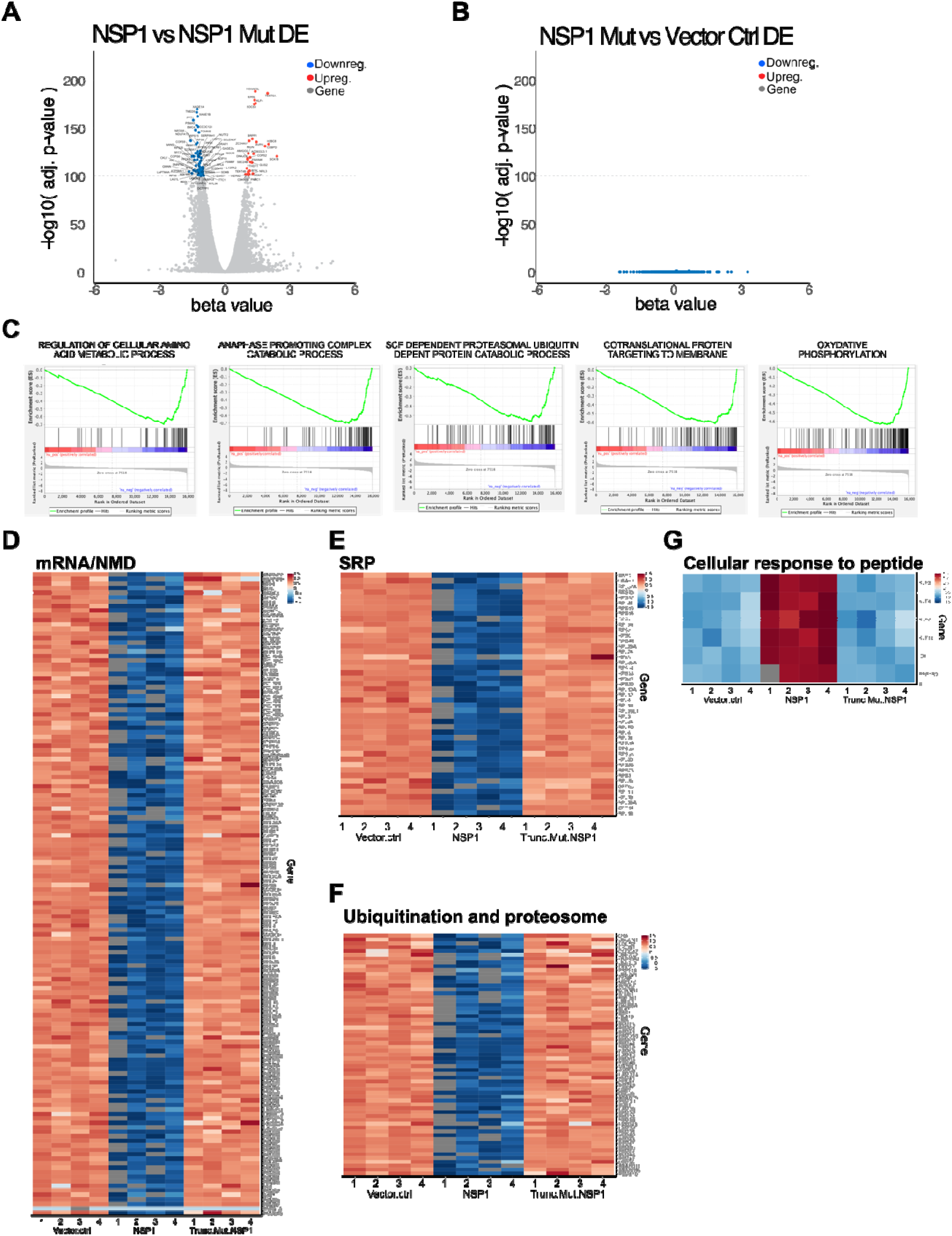
Additional differential expression and pathway analysis of H1299 Nsp1 mRNA-seq dataset, Related to Figures 2 and 3. **(A)** Volcano plot of differential expression between of Nsp1 vs Nsp1 mutant electroporated cells. Genes highly differentially expressed (FDR adjusted q value < 1e-100) are shown with gene names. Upregulated genes are shown in orange. Downregulated genes are shown in blue. **(B)** Volcano plot of differential expression between of Nsp1 mutant vs Vector Control electroporated cells. As seen in the plot, no gene in the genome is differentially expressed between these two groups. **(C)** Gene set enrichment plots of additional representative enriched pathways by GSEA. **(D)** Heatmap of Nsp1 highly repressed genes (q < 1e-30) in the mRNA processing and nonsense-mediated decay processes. **(E)** Heatmap of Nsp1 highly repressed genes (q < 1e-30) in the SRP proteins. **(F)** Heatmap of Nsp1 highly repressed genes (q < 1e-30) in the ubiquitination and proteasome degradation processes. **(G)** Heatmap of Nsp1 highly induced genes (q < 1e-30) in the cellular response to peptide processes.

**Figure S3.**
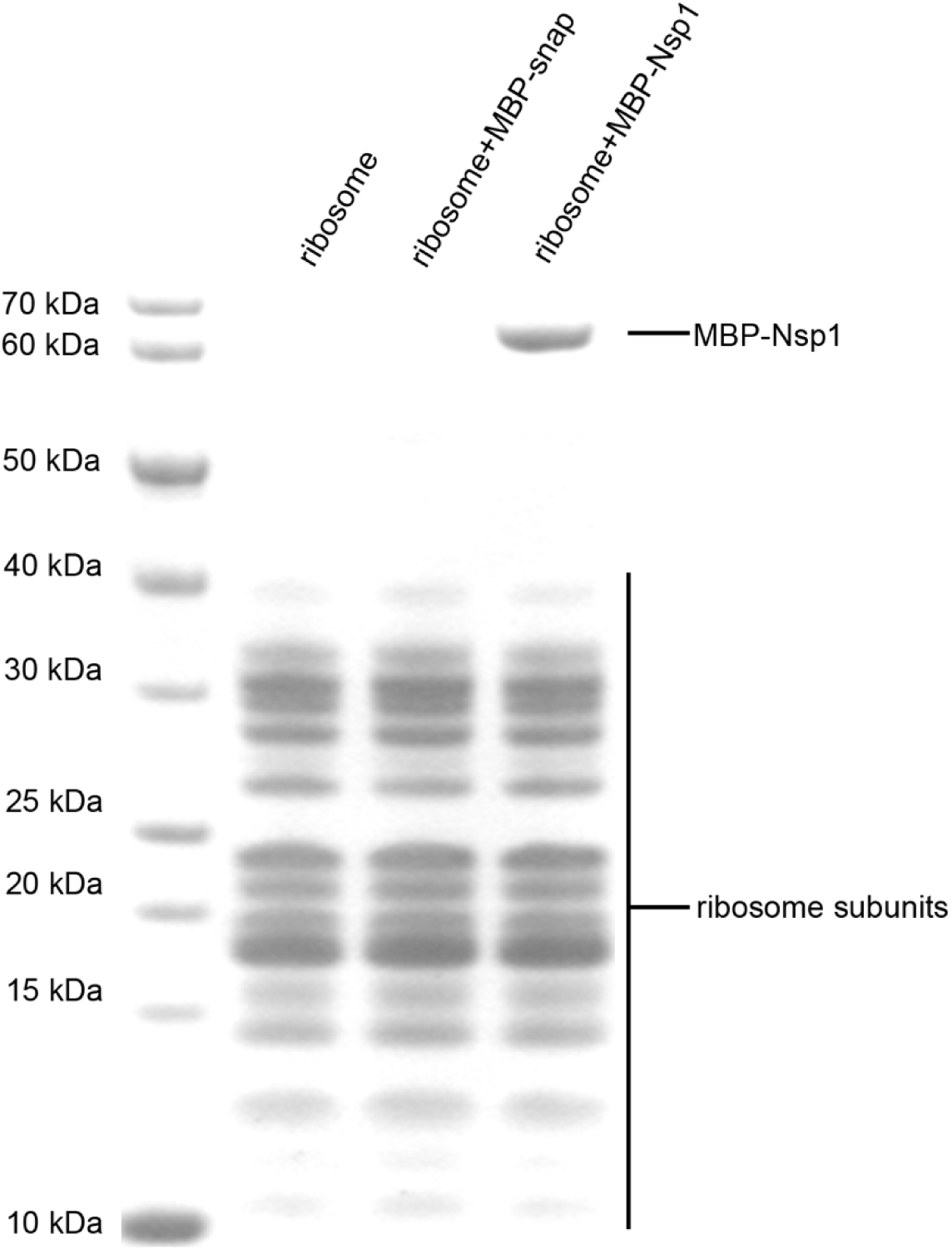
SDS-PAGE analysis of Nsp1 and 40S ribosome binding, Related to Figure 4. Nsp1 is labeled with an MBP tag. MBP-snap was used as a negative control.

**Figure S4.**
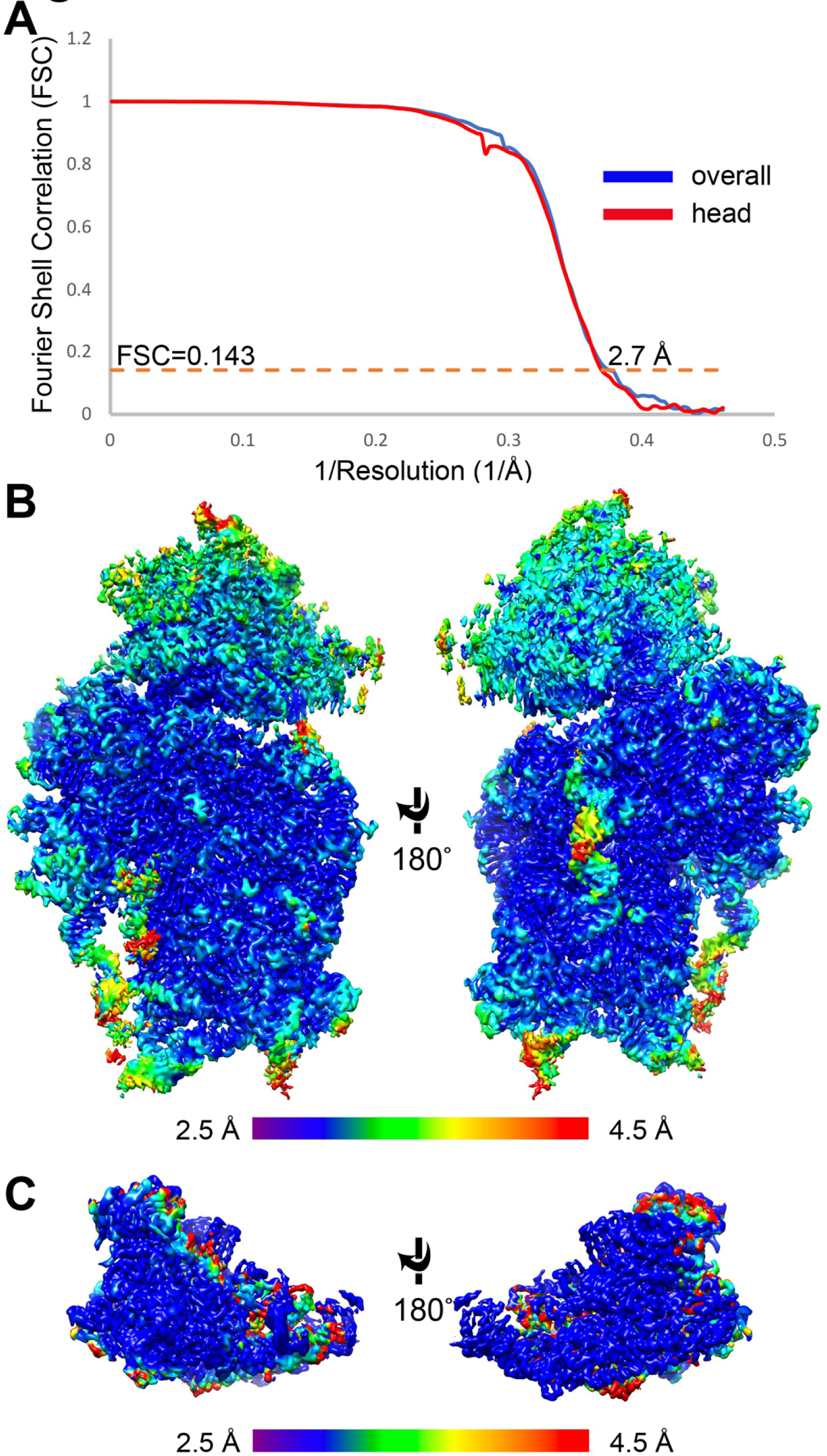
Data processing of Nsp1-40S ribosome complex cryo-EM dataset, Related to Figure 4. **(A)** FSC curves of the half-maps from gold standard refinement of the Nsp1-40S ribosome complex (blue) and masked local refinement of the head domain (red). **(B-C)** Color coded local resolution estimation of the overall complex (**B**) and local-refined head domain (**C**).

**Figure S5.**
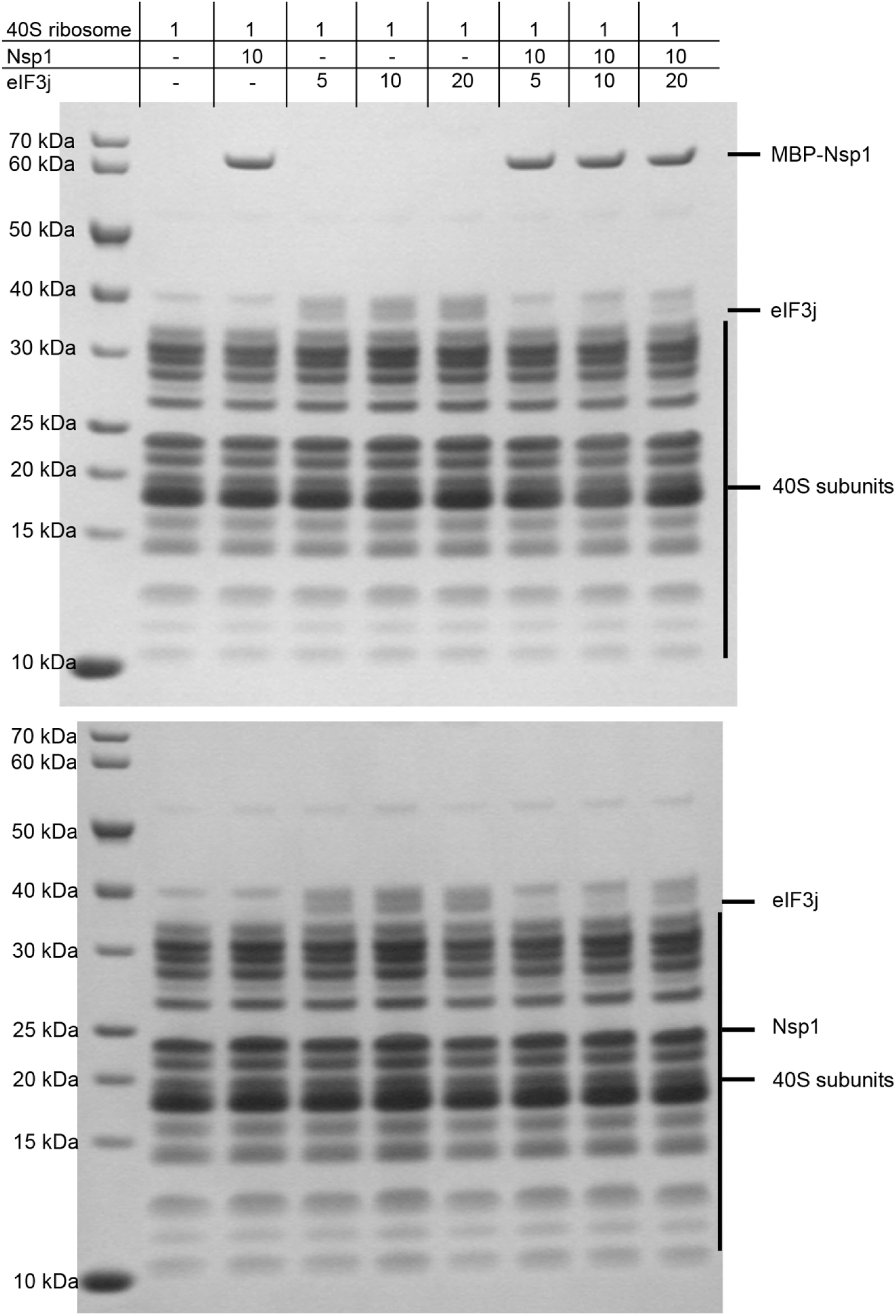
SDS-PAGE analysis of Nsp1 and eIF3j competition assay, Related to Figure 6. Concentration ratios are shown in top table. Top gel: Assay with MBP-Nsp1. Bottom gel: Full-length Nsp1 without the MBP tag was used to exclude the tag effect.

**Figure S6.**
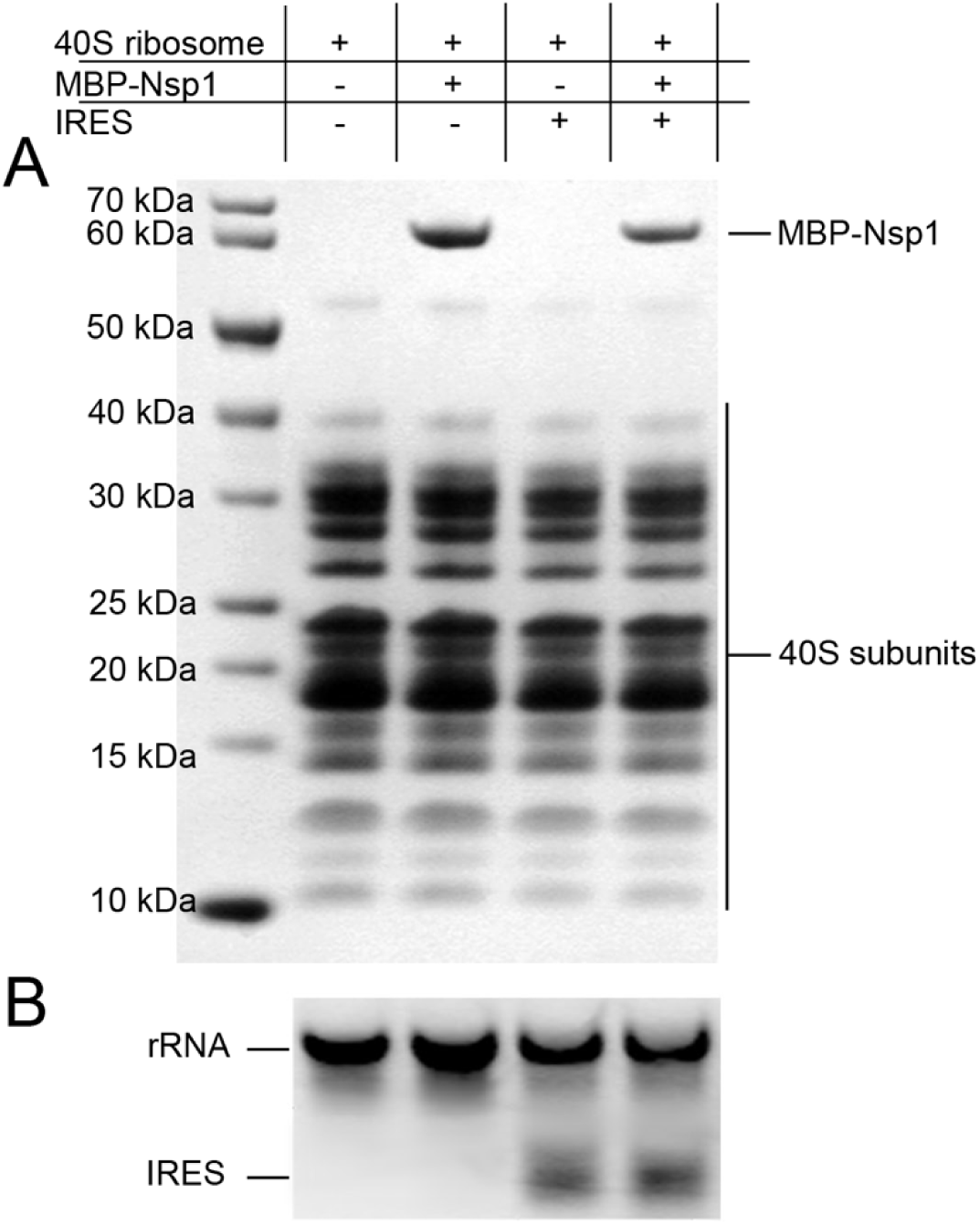
CrPV IRES and Nsp1 can bind to 40S ribosome simultaneously, Related to Figure 6. SDS-PAGE analysis **(A)** and RNA gel **(B)** show the binding of Nsp1 and CrPV IRES.

**Figure S7.**
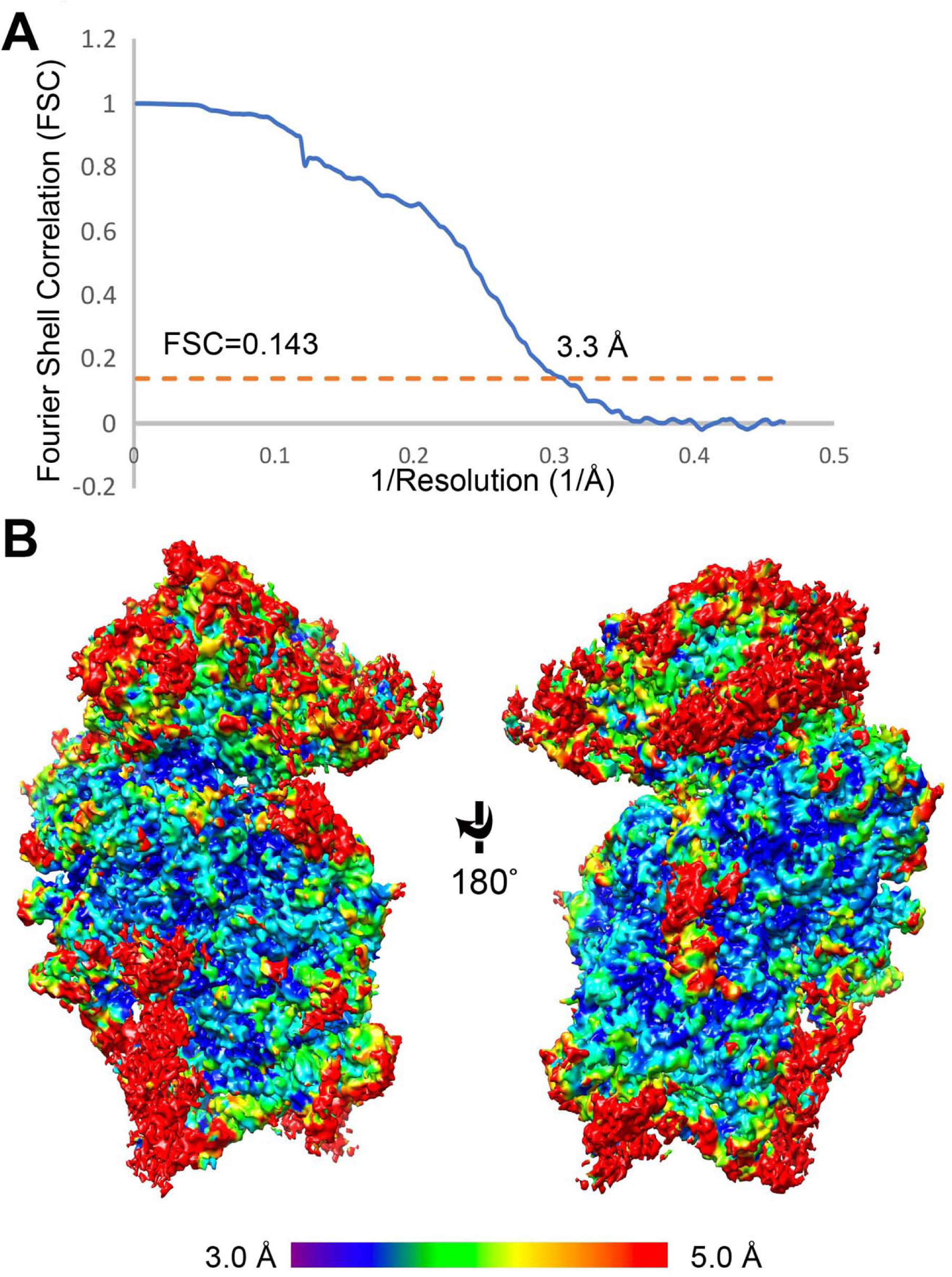
Data processing of Nsp1-40S-CrPV IRES complex cryo-EM dataset, Related to Figure 6. **(A)** FSC curves of the half-maps from gold standard refinement of the Nsp1-40S-CrPV IRES complex. **(B)** Color coded local resolution estimation of the complex.

## List of Supplemental Tables (provided as excel files)

**Table S1. Oligo sequences used in this study**

**Table S2. Source data and summary statistics of cellular viability effect by introduction of SARS-CoV-2 viral proteins and mutants**

**Table S3. Processed Nsp1 mRNA-seq dataset and differential expression analysis**

Sup table 3.1 TPM table of Nsp1 mRNA-seq dataset

Sup table 3.2 Differential expression Nsp1 vs Vector Control

Sup table 3.3 Differential expression Nsp1 Mutant vs Vector Control

Sup table 3.4 Differential expression Nsp1 vs Nsp1 Mutant

**Table S4. DAVID pathway analysis of Nsp1 differentially expressed gene sets**

Sup table 4.1 Functional clustering of Nsp1 vs Vector Control highly downregulated genes (q < 1e-30)

Sup table 4.2 Functional clustering of Nsp1 vs Nsp1 Mutant highly downregulated genes (q < 1e-30)

Sup table 4.3 Functional clustering of Nsp1 vs Vector Control highly upregulated genes (q < 1e-30)

Sup table 4.4 Functional clustering of Nsp1 vs Nsp1 Mutant highly upregulated genes (q < 1e-30)

Sup table 4.5 Biological processes enrichment of Nsp1 vs Vector Control highly downregulated genes (q < 1e-30)

Sup table 4.6 Biological processes enrichment of Nsp1 vs Nsp1 Mutant highly downregulated genes (q < 1e-30)

Sup table 4.7 Biological processes enrichment of Nsp1 vs Vector Control highly upregulated genes (q < 1e-30)

Sup table 4.8 Biological processes enrichment of Nsp1 vs Nsp1 Mutant highly upregulated genes (q < 1e-30)

Sup table 4.9 Gene list of Nsp1 vs Vector Control highly downregulated genes (q < 1e-30)

Sup table 4.10 Gene list enrichment of Nsp1 vs Nsp1 Mutant highly downregulated genes (q < 1e-30)

Sup table 4.11 Gene list enrichment of Nsp1 vs Vector Control highly upregulated genes (q < 1e-30)

Sup table 4.12 Gene list enrichment of Nsp1 vs Nsp1 Mutant highly upregulated genes (q < 1e-30)

Sup table 4.13 Gene list of Nsp1 vs Vector Control all downregulated genes (q < 0.01)

Sup table 4.14 Gene list of Nsp1 vs Vector Control all upregulated genes (q < 0.01)

## STAR Methods

### SARS-CoV-2 plasmid cloning

The cDNA templates of SARS-CoV-2 ORF gene containing plasmids were provided by Dr. Krogan as a gift (Gordon et al., 2020), where the ORFs were primarily cloned into lentiviral expression vector. A non-viral expression vector, pVPSB empty, where ORFs were driven by a constitutive EFS promoter and terminated by a short poly A, was constructed by cloning gBlock fragments (IDT) into pcDNA3.1 vector (Addgene, #52535) by the Gibson assembly (NEB). All ORFs gene encoding fragments were PCR amplified from the lentiviral vectors with ORF-specific forward primers and common reverse primer that containing overlaps that corresponded to flanking sequences of the and KpnI and XhoI restriction sites in the pVPSB empty vector. The primer lists were provided in Table S1. ORFs PCR amplified fragments were gel-purified and cloned into restriction enzyme digested backbone by the Gibson assembly (NEB). A lentiviral vector constitutively expressing a Firefly Luciferase and a puromycin mammalian selection marker (Lenti-Fluc-Puro) was generated by standard molecular cloning. All plasmids were sequenced and harvested by Maxiprep for following assay.

### Nsp1 mutant ORF construction

Truncation mutant Nsp1 has triple stop codons introduced after residues 12 (N terminal mutant). Nsp1 mutant3 has R124 and K125 replaced with S124 and E125 (R124S/K125E). Nsp1 mutant4 has N128 and K129 were converted to S128 and E129 (N128S/K129E). IDT gBlocks were ordered for truncated Nsp1 and different Nsp1 mutants with 19∼23 bp overlaps that corresponded to flanking sequences of the and AgeI and BstXI restriction sites in the pVPSBA01-Nsp1 plasmid. pVPSBA01-Nsp1 plasmid were digested and gel purified, and gBlocks were cloned using the Gibson assembly (NEB).

### Generation of stable cell lines

Lentivirus was produced by transfection of co-transgene plasmid (Lenti-Fluc-Puro) and packaging plasmids (psPAX2, pMD2.G) into HEK293FT cells, followed by supernatant harvesting, filtering and concentration with Amicon filters (Sigma). H1299 and Vero E6 cells were infected with Lenti-Fluc-Puro lentivirus. After 24 h of virus transduction, cells were selected with 10 μg/mL puromycin, until all cells died in the control group. Luc expressing H1299 and Vero E6 that with puromycin resistance cell lines were obtained and named as H1299-PL and Vero E6-PL (Vero E6-PL for short) respectively.

### Mammalian cell culture

H1299, H1299-PL, Vero E6, Vero E6-PL cell lines were cultured in Dulbecco’s modified Eagle’ s medium (DMEM; Thermo fisher) supplemented with 10% Fetal bovine serum (FBS, Hyclone),1% penicillin-streptomycin (Gibco), named as D10 medium. Cells were typically passaged every 1-2 days at a split ratio of 1:2 or 1:4 when the confluency reached at 80%.

### SARS-CoV-2 ORF mini-screen for cell viability

H1299 cells were plated in white opaque walled microwell assay plates, 25,000 cells per 96 well. SARS-CoV-2 ORF plasmids, 1 μg of each, were parallelly transfected with 1 μl lipofectamine 2000, in triplicates. Cell viability was detected at every 24hr after transfection using CellTiter-Glo® Luminescent Cell Viability Assay kit (Promega). Relative viability was normalized to the mean viability of empty vector transfected control group. All procedures followed the manufacturer standard protocol. Luminescent signals were measured by a Plate Reader (PerkinElmer).

### Determination of luciferase reporter cell viability

H1299-PL and Vero E6-PL cells were plated in white opaque walled microwell assay plates, 25,000 cells per well in a 96 well. SARS-CoV-2 ORF plasmids, 1 μg of each, were parallelly transfected with 1ul lipofectamine 2000. Cell viability was measured every 24 hr after plasmid transfection by adding 150 μg / ml D-Luciferin (PerkinElmer) using a multi-channel pipette. Luciferase intensity was measured by a Plate Reader (PerkinElmer).

### Electroporation with 4D nucleofection

Cells were trypsinized and collected, 1e6 cells were resuspended in SF cell line NucleofectorTM solution with 3 μg plasmid DNA. Cells were transferred into 100 μl NucleocuvetteTM Vessel and NCI-H1299 [H1299] cell specific protocol were utilized according to the manufacturer’s protocol (4D-NucleofectorTM X Unit, Lonza). After the pulse application, 100 μl prewarmed D10 medium was added to the electroporated cells in the cuvette. Cells were gently resuspended in the cuvette and transferred into 6 well plate, cultured in incubator. Cells were collected at 24 or 48 hours later for flowcytometry assay and RNA extraction.

### Apoptosis flow cytometry assay

Flow cytometry was performed using standard immunology protocols. Briefly, experimental and control cells were electroporated with respective plasmids. After a defined time point, cells were collected, fixed and permeabilized using Fixation/Permeablization Solution kit (BD). Then antigen-specific antibodies with specific dilutions were added into cells and incubated for 30 min on ice. Cells were washed with cold MACS buffer for 3 times before analyzed on a BD FACSAria cytometer. Antibody used: anti-cleaved Caspase-3(Asp175) (Sigma, 9669s, 1:200).

### Gene expression analysis by mRNA sequencing (mRNA-seq, RNA-seq)

For H1299-PL cells electroporated with Nsp1 or Nsp1 mutant, mRNA-seq libraries were prepared following next-generation sequencing (NGS) protocols. Briefly, 1e6 H1299 cells were electroporated with 3 μg Nsp1, mutant Nsp1, and relative control plasmids. Electroporation was done in with quadruplicates for each group. Cells were collected 24hr post electroporation. Total mRNA was extracted with RNasy Plus Mini Kit (Qiagen). 1μg total mRNA each sample was used for the RNA-seq library preparations. A NEBNext® Ultra™ RNA Library Prep Kit for Illumina was employed to perform RNA-seq library preparation and samples were multiplexed using barcoded primers provided by NEBNext® Multiplex Oligos for Illumina® (Index Primers Set 1). All procedures follow the manufacturer standard protocol. Libraries were sequenced with Novaseq system (Illumina).

### mRNA-seq data processing, differential expression analysis and pathway analysis

The mRNA data processing, transcript quantification, differential expression, and pathway analysis were performed using custom computational programs. In brief, Fastq files from mRNA-seq were used analyzed using the Kallisto quant algorithm for transcript quantification (Bray et al., 2016). Differential expression analysis was performed using Sleuth (Pimentel et al., 2017). Z-scores for time course heatmap were calculated by log2-normalizion of gene counts following by scaling by genes. Visualizations of differentially expressed genes such as volcano plots and heatmaps were generated using standard R packages. Differentially upregulated and downregulated genes were subjected to pathway analysis by DAVID (Huang et al., 2007) and/or GSEA (Subramanian et al., 2005). Processed mRNA-seq data, differential expression analysis and pathway analysis results are provided in (Table S3 and S4).

### RT-qPCR

Total RNA was extracted from cells using RNasy Plus Mini Kit (Qiagen). Total mRNA was reverse transcribed into cDNA by M-MLV Reverse Transcriptase (Sigma). Samples were collected in triplicates. Gene expression was quantified using Taqman Fast Universal PCR Master Mix (Thermo Fisher) and Taqman probes (Invitrogen). NSP1 probe was generated with custom designed according to the Nsp1 DNA sequence in the SARS-CoV-2 genome annotation (2019-nCoV/USA-WA1/2020, accession MN985325). RNA expression level was normalized to *ACTB* (human). Relative mRNA expression was determined via the ΔΔ *C*_t_ method.

### Sample size determination

Sample size was determined according to the lab’s prior work or similar approaches in the field.

### Replication

All experiments were done with at least three biological replicates. Experimental replications were indicated in detail in methods section and in each figure panel’s legend.

### Standard statistical analysis

All statistical methods are described in figure legends and/or supplementary Excel tables. The P values and statistical significance were estimated for all analyses. For example, the unpaired, two-sided, T test was used to compare two groups. One-way ANOVA along with multiple comparisons test, was used to compare multiple groups. Multiple-testing correction was done using false discovery rate (FDR) method. Different levels of statistical significance were accessed based on specific p values and type I error cutoffs (0.05, 0.01, 0.001, 0.0001). Data analysis was performed using GraphPad Prism v.8. and/or RStudio.

### Ribosome and CrPV IRES purification

40S ribosomal subunits were purified from the rabbit reticulocyte lysate (Green Hectares, USA) as described previously (Lomakin and Steitz, 2013). The gene for wild-type CrPV IRES (nucleotides 6028-6240) was chemically synthesized and cloned in the pBluescript SK vector flanked at the 5’-end by a T7 promoter sequence and an EcoRI cleavage site at the 3’-end. Standard *in vitro* transcription protocol was used for IRES RNA synthesis and purification (MEGAscript™ T7 Transcription Kit, Ambion, USA).

### Protein construction, expression and purification

Full-length SARS-CoV-2 Nsp1 was cloned into pMAT-9s vector and pET-Duet vector for expression of MBP-tagged and 6×his tagged proteins, respectively. The Escherichia coli BL21 (DE3) cells were used for protein expressions, which were induced by 0.5 mM isopropyl β-D-1-thiogalactopyranoside (IPTG) at 16 °C for 16 hours in Terrific Broth. Cells were harvested and lysed using a microfluidizer. The lysate was clarified by centrifugation and then applied to a Ni-NTA (Qiagen) column. Anion exchange (HiTrap Q HP, GE healthcare) chromatography was performed in a buffer of 50 mM Tris, pH 8.0 with a NaCl concentration gradient from 50 mM to 1M. Subsequent size exclusion chromatography (HiLoad Superdex 75, GE healthcare) was performed in a buffer of 50 mM Tris, 150 mM NaCl, pH 8.0. Purity of the proteins was analyzed by SDS-PAGE after each step. Full length eIF3j was expressed in Escherichia coli BL21 and purified with a similar method.

### Filter binding assays

Rabbit 40S ribosome and binding partners (proteins or CrPV IRES RNA) were incubated together for 20 min at 37 °C in a total volume of 20 μl in 1× 48S buffer (20 mM HEPES(KOH) pH 7.5, 100 mM KCl, 2.5 mM MgAc, 1 mM DTT, 250 μM Spermidine 3HCl). Reaction mixtures were incubated for another 20 min at room temperature before diluting to 100 μl with H100 buffer (10 mM HEPES(KOH) pH 7.0, 100 mM KCl, 5 mM MgAc, 2 mM DTT). Diluted reaction mixtures were filtered through 100 kDa filter (Thermo Scientific) in 10,000g for 5 min. The flow through was collected. 200 μl H100 buffer was used for washing the unbound proteins or RNA for 4 times before analyzing by SDS-PAGE or RNA gel.

The concentration for the 40S ribosome for the filter binding assay is 1.5 μM and the Nsp1 concentration is 15 μM (ratio of 1:10). In the Nsp1 and eIF3j competition assays, the concentrations of eIF3j are 7.5 μM, 15 μM and 30 μM corresponding to ratios of 1:5, 1:10 and 1:20. The concentration of the CrPV IRES is 7.5 μM in the Nsp1-IRES binding assay (ratio of 1:5).

### Cryo-EM sample preparation, data collection and processing

40S ribosome and Nsp1, with or without the CrPV IRES RNA were mixed and incubated at 37 °C for 20 mins to form a stable complex. The complex (4 μl) was applied to a C899 Flat 2/1 3C copper grid (Electron Microscopy Sciences) pretreated by glow-discharging at 8 mA for 20 seconds. The grid was blotted at 20 °C with 100% humidity and plunge-frozen in liquid ethane using FEI Vitrobot Mark IV (Thermo Fisher). The grids were stored in liquid nitrogen before data collection.

Images were acquired on a FEI Titan Krios electron microscope (Thermo Fisher) equipped with a post-GIF Gatan K3 direct detector in super-resolution mode, at a nominal calibrated magnification of 81,000× with the physical pixel size corresponding to 1.068Å. Automated data collection was performed using SerialEM (Mastronarde, 2005).

A total of 4,700 movie series were collected for the Nsp1-40S ribosome complex. 300 movies series were collected for the Nsp1-40S-CrPV IRES complex. For the Nsp1-40S ribosome complex, a defocus range of 0.5 μm to 2 μm was used. Data were collected with a dose of 15.9 electrons per pixel per second. Images were recorded over a 3.6s exposure with 0.1s for each frame to give a total dose of 50 electrons per Å^2^. Similar conditions were used for the Nsp1-40S-CrPV IRES complex.

The same data processing procedures were carried out for both the two complexes using standard pipelines in cryoSPARC(Punjani et al., 2017). The final average resolution is 2.7 Å for the Nsp1-40S ribosome complex and 3.3 Å for the Nsp1-40S-CrPV IRES complex (FSC=0.143). Local refinement was carried out for the head domain of the 40S, which significantly increased the quality of the reconstruction for this domain (Figure S4C).

### Model building and refinement

The structure of the rabbit 40S ribosome was extracted from PDB: 4KZX (Lomakin and Steitz, 2013) and 6SGC (Chandrasekaran et al., 2019). The model of Nsp1 C-terminal domain was manually built in COOT (Emsley et al., 2010). The CrPV IRES structure was extracted form PDB:5IT9 and refined (Murray et al., 2016). The structures of Nsp1-40S ribosome complex and Nsp1-IRES-40S ribosome complex were refined with phenix.real_space_refine module in PHENIX (Adams et al., 2010). All structural figures were generated using PyMol (Schrodinger, 2015) and Chimera (Pettersen et al., 2004).

## Data and resource availability

All data generated or analyzed during this study are included in this article and its supplementary information files. Specifically, source data and statistics for non-high-throughput experiments are provided in a supplementary table excel file (Table S2). High-throughput experiment data are provided as processed quantifications in Supplemental Datasets (Table S3 and S4). Genomic sequencing raw data are being deposited to NIH Sequence Read Archive (SRA) and/or Gene Expression Omnibus (GEO), with pending accession numbers. Constructs are available at either through a public repository or via requests to the corresponding authors. Original cell lines are available at commercial sources listed in supplementary information files. Genetically modified cell lines are available via the authors’ laboratories. Codes that support the findings of this research are being deposited to a public repository such as GitHub, and are available from the corresponding authors upon reasonable request.

The cryo-EM maps of the Nsp1-40S ribosome complex and the Nsp1-40S-CrPV IRES ribosome complex have been deposited in the Electron Microscopy Data Bank as EMD-XXXX and EMD-YYYY, respectively. The corresponding structure models are in the Protein Data Bank with accession code XXXX, YYYY.

## References

Adams, P.D., Afonine, P.V., Bunkoczi, G., Chen, V.B., Davis, I.W., Echols, N., Headd, J.J., Hung, L.W., Kapral, G.J., Grosse-Kunstleve, R.W., et al. (2010). PHENIX: a comprehensive Python-based system for macromolecular structure solution. Acta Crystallogr D Biol Crystallogr 66, 213–221.

Almeida, M.S., Johnson, M.A., Herrmann, T., Geralt, M., and Wuthrich, K. (2007). Novel beta-barrel fold in the nuclear magnetic resonance structure of the replicase nonstructural protein 1 from the severe acute respiratory syndrome coronavirus. J Virol 81, 3151–3161.

Astuti, I., and Ysrafil (2020). Severe Acute Respiratory Syndrome Coronavirus 2 (SARS-CoV-2): An overview of viral structure and host response. Diabetes Metab Syndr 14, 407–412.

Aylett, C.H., Boehringer, D., Erzberger, J.P., Schaefer, T., and Ban, N. (2015). Structure of a yeast 40S-eIF1-eIF1A-eIF3-eIF3j initiation complex. Nat Struct Mol Biol 22, 269–271.

Babaylova, E., Malygin, A., Gopanenko, A., Graifer, D., and Karpova, G. (2019a). Tetrapeptide 60-63 of human ribosomal protein uS3 is crucial for translation initiation. Biochim Biophys Acta Gene Regul Mech 1862, 194411.

Babaylova, E., Malygin, A., Gopanenko, A., Graifer, D., and Karpova, G. (2019b). Tetrapeptide 60-63 of human ribosomal protein uS3 is crucial for translation initiation. Bba-Gene Regul Mech 1862.

Blanco-Melo, D., Nilsson-Payant, B.E., Liu, W.C., Uhl, S., Hoagland, D., Moller, R., Jordan, T.X., Oishi, K., Panis, M., Sachs, D., et al. (2020). Imbalanced Host Response to SARS-CoV-2 Drives Development of COVID-19. Cell 181, 1036–1045 e1039.

Bray, N.L., Pimentel, H., Melsted, P., and Pachter, L. (2016). Near-optimal probabilistic RNA-seq quantification. Nat Biotechnol 34, 525–527.

Cate, J.H. (2017). Human eIF3: from ‘blobology’ to biological insight. Philos Trans R Soc Lond B Biol Sci 372.

Chandrasekaran, V., Juszkiewicz, S., Choi, J., Puglisi, J.D., Brown, A., Shao, S., Ramakrishnan, V., and Hegde, R.S. (2019). Mechanism of ribosome stalling during translation of a poly(A) tail. Nat Struct Mol Biol 26, 1132–1140.

Coronaviridae Study Group of the International Committee on Taxonomy of, V. (2020). The species Severe acute respiratory syndrome-related coronavirus: classifying 2019-nCoV and naming it SARS-CoV-2. Nat Microbiol 5, 536–544.

Dong, J.S., Aitken, C.E., Thakur, A., Shin, B.S., Lorsch, J.R., and Hinnebusch, A.G. (2017). Rps3/uS3 promotes mRNA binding at the 40S ribosome entry channel and stabilizes preinitiation complexes at start codons. P Natl Acad Sci USA 114, E2126–E2135.

Emsley, P., Lohkamp, B., Scott, W.G., and Cowtan, K. (2010). Features and development of Coot. Acta Crystallogr D Biol Crystallogr 66, 486–501.

Fraser, C.S., Berry, K.E., Hershey, J.W.B., and Doudna, J.A. (2007). eIF3j is located in the decoding center of the human 40S ribosomal subunit. Mol Cell 26, 811–819.

Fraser, C.S., Lee, J.Y., Mayeur, G.L., Bushell, M., Doudna, J.A., and Hershey, J.W. (2004a). The j-subunit of human translation initiation factor eIF3 is required for the stable binding of eIF3 and its subcomplexes to 40 S ribosomal subunits in vitro. J Biol Chem 279, 8946–8956.

Fraser, C.S., Lee, J.Y., Mayeur, G.L., Bushell, M., Doudna, J.A., and Hershey, J.W.B. (2004b). The j-subunit of human translation initiation factor eIF3 is required for the stable binding of eIF3 and its subcomplexes to 40 S ribosomal subunits in vitro. J Biol Chem 279, 8946–8956.

Gordon, D.E., Jang, G.M., Bouhaddou, M., Xu, J., Obernier, K., White, K.M., O’Meara, M.J., Rezelj, V.V., Guo, J.Z., Swaney, D.L., et al. (2020). A SARS-CoV-2 protein interaction map reveals targets for drug repurposing. Nature 583, 459–468.

Graifer, D., Malygin, A., Zharkov, D.O., and Karpova, G. (2014). Eukaryotic ribosomal protein S3: A constituent of translational machinery and an extraribosomal player in various cellular processes. Biochimie 99, 8–18.

Grasselli, G., Zangrillo, A., Zanella, A., Antonelli, M., Cabrini, L., Castelli, A., Cereda, D., Coluccello, A., Foti, G., Fumagalli, R., et al. (2020). Baseline Characteristics and Outcomes of 1591 Patients Infected With SARS-CoV-2 Admitted to ICUs of the Lombardy Region, Italy. JAMA.

Haimov, O., Sinvani, H., Martin, F., Ulitsky, I., Emmanuel, R., Tamarkin-Ben-Harush, A., Vardy, A., and Dikstein, R. (2017). Efficient and Accurate Translation Initiation Directed by TISU Involves RPS3 and RPS10e Binding and Differential Eukaryotic Initiation Factor 1A Regulation. Mol Cell Biol 37.

Hashem, Y., des Georges, A., Dhote, V., Langlois, R., Liao, H.Y., Grassucci, R.A., Pestova, T.V., Hellen, C.U., and Frank, J. (2013). Hepatitis-C-virus-like internal ribosome entry sites displace eIF3 to gain access to the 40S subunit. Nature 503, 539–543.

Hershey, J.W. (2015). The role of eIF3 and its individual subunits in cancer. Biochim Biophys Acta 1849, 792–800.

Hinnebusch, A.G. (2014). The scanning mechanism of eukaryotic translation initiation. Annu Rev Biochem 83, 779–812.

Hinnebusch, A.G. (2017a). Structural Insights into the Mechanism of Scanning and Start Codon Recognition in Eukaryotic Translation Initiation. Trends Biochem Sci 42, 589–611.

Hinnebusch, A.G. (2017b). Structural Insights into the Mechanism of Scanning and Start Codon Recognition in Eukaryotic Translation Initiation. Trends Biochem Sci 42, 589–611.

Hinnebusch, A.G., Ivanov, I.P., and Sonenberg, N. (2016). Translational control by 5’-untranslated regions of eukaryotic mRNAs. Science 352, 1413–1416.

Hoffmann, M., Kleine-Weber, H., Schroeder, S., Kruger, N., Herrler, T., Erichsen, S., Schiergens, T.S., Herrler, G., Wu, N.H., Nitsche, A., et al. (2020). SARS-CoV-2 Cell Entry Depends on ACE2 and TMPRSS2 and Is Blocked by a Clinically Proven Protease Inhibitor. Cell 181, 271-+.

Huang, C., Lokugamage, K.G., Rozovics, J.M., Narayanan, K., Semler, B.L., and Makino, S. (2011). SARS coronavirus nsp1 protein induces template-dependent endonucleolytic cleavage of mRNAs: viral mRNAs are resistant to nsp1-induced RNA cleavage. Plos Pathog 7, e1002433.

Huang, D.W., Sherman, B.T., Tan, Q., Collins, J.R., Alvord, W.G., Roayaei, J., Stephens, R., Baseler, M.W., Lane, H.C., and Lempicki, R.A. (2007). The DAVID Gene Functional Classification Tool: a novel biological module-centric algorithm to functionally analyze large gene lists. Genome Biol 8, R183.

Jan, E., Mohr, I., and Walsh, D. (2016). A Cap-to-Tail Guide to mRNA Translation Strategies in Virus-Infected Cells. Annu Rev Virol 3, 283–307.

Kamitani, W., Huang, C., Narayanan, K., Lokugamage, K.G., and Makino, S. (2009). A two-pronged strategy to suppress host protein synthesis by SARS coronavirus Nsp1 protein. Nat Struct Mol Biol 16, 1134–U1132.

Kamitani, W., Narayanan, K., Huang, C., Lokugamage, K., Ikegami, T., Ito, N., Kubo, H., and Makino, S. (2006). Severe acute respiratory syndrome coronavirus nsp1 protein suppresses host gene expression by promoting host mRNA degradation. Proc Natl Acad Sci U S A 103, 12885–12890.

Kim, D., Lee, J.Y., Yang, J.S., Kim, J.W., Kim, V.N., and Chang, H. (2020). The Architecture of SARS-CoV-2 Transcriptome. Cell 181, 914–921 e910.

Lee, A.S., Kranzusch, P.J., Doudna, J.A., and Cate, J.H. (2016). eIF3d is an mRNA cap-binding protein that is required for specialized translation initiation. Nature 536, 96–99.

Lim, Y.X., Ng, Y.L., Tam, J.P., and Liu, D.X. (2016). Human Coronaviruses: A Review of Virus-Host Interactions. Diseases 4.

Lokugamage, K.G., Narayanan, K., Huang, C., and Makino, S. (2012). Severe Acute Respiratory Syndrome Coronavirus Protein nsp1 Is a Novel Eukaryotic Translation Inhibitor That Represses Multiple Steps of Translation Initiation. J Virol 86, 13598–13608.

Lomakin, I.B., and Steitz, T.A. (2013). The initiation of mammalian protein synthesis and mRNA scanning mechanism. Nature 500, 307–311.

Lozano, G., and Martinez-Salas, E. (2015). Structural insights into viral IRES-dependent translation mechanisms. Curr Opin Virol 12, 113–120.

Martinez-Salas, E., Francisco-Velilla, R., Fernandez-Chamorro, J., and Embarek, A.M. (2018). Insights into Structural and Mechanistic Features of Viral IRES Elements. Front Microbiol 8.

Masters, P.S. (2006). The molecular biology of coronaviruses. Adv Virus Res 66, 193–292.

Mastronarde, D.N. (2005). Automated electron microscope tomography using robust prediction of specimen movements. J Struct Biol 152, 36–51.

Murray, J., Savva, C.G., Shin, B.S., Dever, T.E., Ramakrishnan, V., and Fernandez, I.S. (2016). Structural characterization of ribosome recruitment and translocation by type IV IRES. Elife 5.

Narayanan, K., Huang, C., Lokugamage, K., Kamitani, W., Ikegami, T., Tseng, C.T., and Makino, S. (2008). Severe acute respiratory syndrome coronavirus nsp1 suppresses host gene expression, including that of type I interferon, in infected cells. J Virol 82, 4471–4479.

Niepmann, M., and Gerresheim, G.K. (2020). Hepatitis C Virus Translation Regulation. Int J Mol Sci 21.

Passmore, L.A., Schmeing, T.M., Maag, D., Applefield, D.J., Acker, M.G., Algire, M.A., Lorsch, J.R., and Ramakrishnan, V. (2007). The eukaryotic translation initiation factors eIF1 and eIF1A induce an open conformation of the 40S ribosome. Mol Cell 26, 41–50.

Pettersen, E.F., Goddard, T.D., Huang, C.C., Couch, G.S., Greenblatt, D.M., Meng, E.C., and Ferrin, T.E. (2004). UCSF Chimera--a visualization system for exploratory research and analysis. J Comput Chem 25, 1605–1612.

Pimentel, H., Bray, N.L., Puente, S., Melsted, P., and Pachter, L. (2017). Differential analysis of RNA-seq incorporating quantification uncertainty. Nat Methods 14, 687–690.

Pisarev, A.V., Kolupaeva, V.G., Yusupov, M.M., Hellen, C.U., and Pestova, T.V. (2008). Ribosomal position and contacts of mRNA in eukaryotic translation initiation complexes. The EMBO journal 27, 1609–1621.

Punjani, A., Rubinstein, J.L., Fleet, D.J., and Brubaker, M.A. (2017). cryoSPARC: algorithms for rapid unsupervised cryo-EM structure determination. Nat Methods 14, 290–296.

Schrodinger, LLC (2015). The PyMOL Molecular Graphics System, Version 1.8.

Sharifulin, D.E., Bartuli, Y.S., Meschaninova, M.I., Ven’yaminova, A.G., Graifer, D.M., and Karpova, G.G. (2016). Exploring accessibility of structural elements of the mammalian 40S ribosomal mRNA entry channel at various steps of translation initiation. Bba-Proteins Proteom 1864, 1328–1338.

Sharifulin, D.E., Grosheva, A.S., Bartuli, Y.S., Malygin, A.A., Meschaninova, M.I., Ven’yaminova, A.G., Stahl, J., Graifer, D.M., and Karpova, G.G. (2015). Molecular contacts of ribose-phosphate backbone of mRNA with human ribosome. Bba-Gene Regul Mech 1849, 930–939.

Sokabe, M., and Fraser, C.S. (2014). Human eukaryotic initiation factor 2 (eIF2)-GTP-Met-tRNAi ternary complex and eIF3 stabilize the 43 S preinitiation complex. J Biol Chem 289, 31827–31836.

Subramanian, A., Tamayo, P., Mootha, V.K., Mukherjee, S., Ebert, B.L., Gillette, M.A., Paulovich, A., Pomeroy, S.L., Golub, T.R., Lander, E.S., et al. (2005). Gene set enrichment analysis: a knowledge-based approach for interpreting genome-wide expression profiles. Proc Natl Acad Sci U S A 102, 15545–15550.

Tanaka, T., Kamitani, W., DeDiego, M.L., Enjuanes, L., and Matsuura, Y. (2012). Severe acute respiratory syndrome coronavirus nsp1 facilitates efficient propagation in cells through a specific translational shutoff of host mRNA. J Virol 86, 11128–11137.

Thoms, M., Buschauer, R., Ameismeier, M., Koepke, L., Denk, T., Hirschenberger, M., Kratzat, H., Hayn, M., Mackens-Kiani, T., Cheng, J., et al. (2020). Structural basis for translational shutdown and immune evasion by the Nsp1 protein of SARS-CoV-2. Science.

Tohya, Y., Narayanan, K., Kamitani, W., Huang, C., Lokugamage, K., and Makino, S. (2009). Suppression of host gene expression by nsp1 proteins of group 2 bat coronaviruses. J Virol 83, 5282–5288.

Walker, M.J., Shortridge, M.D., Albin, D.D., Cominsky, L.Y., and Varani, G. (2020). Structure of the RNA Specialized Translation Initiation Element that Recruits eIF3 to the 5’-UTR of c-Jun. J Mol Biol 432, 1841–1855.

Walsh, D., and Mohr, I. (2011). Viral subversion of the host protein synthesis machinery. Nat Rev Microbiol 9, 860–875.

Wathelet, M.G., Orr, M., Frieman, M.B., and Baric, R.S. (2007). Severe acute respiratory syndrome coronavirus evades antiviral signaling: role of nsp1 and rational design of an attenuated strain. J Virol 81, 11620–11633.

Wei, J., Alfajaro, M.M., Hanna, R.E., DeWeirdt, P.C., Strine, M.S., Lu-Culligan, W.J., Zhang, S.-M., Graziano, V.R., Schmitz, C.O., Chen, J.S., et al. (2020). Genome-wide CRISPR screen reveals host genes that regulate SARS-CoV-2 infection. bioRxiv, 2020.2006.2016.155101.

Wong, H.H., Kumar, P., Tay, F.P.L., Moreau, D., Liu, D.X., and Bard, F. (2015). Genome-Wide Screen Reveals Valosin-Containing Protein Requirement for Coronavirus Exit from Endosomes. J Virol 89, 11116–11128.

Yoshimoto, F.K. (2020). The Proteins of Severe Acute Respiratory Syndrome Coronavirus-2 (SARS CoV-2 or n-COV19), the Cause of COVID-19. Protein J 39, 198–216.

Zhou, P., Yang, X.L., Wang, X.G., Hu, B., Zhang, L., Zhang, W., Si, H.R., Zhu, Y., Li, B., Huang, C.L., et al. (2020). A pneumonia outbreak associated with a new coronavirus of probable bat origin. Nature 579, 270–273.

Ziebuhr, J. (2005). The coronavirus replicase. Curr Top Microbiol Immunol 287, 57–94.

Zust, R., Cervantes-Barragan, L., Kuri, T., Blakqori, G., Weber, F., Ludewig, B., and Thiel, V. (2007). Coronavirus non-structural protein 1 is a major pathogenicity factor: Implications for the rational design of coronavirus vaccines. Plos Pathog 3, 1062–1072.

